# Energy metabolism modulates the regulatory impact of activators on gene expression

**DOI:** 10.1101/2023.10.24.563842

**Authors:** Sha Qiao, Sebastian Bernasek, Kevin D. Gallagher, Shigehiro Yamada, Neda Bagheri, Luis A.N. Amaral, Richard W. Carthew

## Abstract

Gene expression is a regulated process fueled by ATP consumption. Therefore, regulation must be coupled to constraints imposed by the level of energy metabolism. Here, we explore this relationship both theoretically and experimentally. A stylized mathematical model predicts that activators of gene expression have variable impact depending on metabolic rate. Activators become less essential when metabolic rate is reduced and more essential when metabolic rate is enhanced. We find that in the *Drosophila* eye, expression dynamics of the *yan* gene are less affected by loss of EGFR-mediated activation when metabolism is reduced, and the opposite effect is seen when metabolism is enhanced. The effects are also seen at the level of pattern regularity in the adult eye, where loss of EGFR-mediated activation is mitigated by lower metabolism. We propose that gene activation is tuned by energy metabolism to allow for faithful expression dynamics in the face of variable metabolic conditions.

## INTRODUCTION

The development of an organism occurs over a period of time that is distinct for the species to which the organism belongs [1]. Since development is coupled to the activities of gene regulatory networks (GRNs) that operate within cells, the dynamical properties of these GRNs are thought to influence the timing of development. For example, the turnover rates of proteins operating within a human neurodevelopmental GRN are slower than for those proteins operating in the homologous mouse GRN, and this difference is thought to partially explain the large difference in developmental tempo between the two species [2, 3].

The pace of development is also dependent upon extrinsic factors such as cellular metabolism [4]. Gene expression requires the continual synthesis of key metabolites and ATP, the primary source of chemical energy. The energy budget of a cell is composed of many competing processes that expend energy by consuming ATP. For example, during embryogenesis, gene expression and cell division account for a small fraction of the energy expended, suggesting that the energy budget is devoted to many biochemical processes [5, 6]. Energy expenditure is balanced with the generation of ATP, which depends on metabolic processing of nutrients. If an organism’s nutrient uptake varies for whatever reason, then energy expenditure must accordingly adjust to prevent exhaustion of ATP stores [7]. This principle was demonstrated in *Drosophila* larvae as they grow and develop. Targeted ablation of insulin-like peptide (Dilp) expression causes larval cells to reduce their uptake of circulating sugars by ∼40% [8]. There is a corresponding 30% decrease in energy expenditure by the body, and as a result, the animals develop more slowly and grow into slightly smaller adults [8]. Developmental gene expression dynamics are correspondingly slower [9].

Since cells adjust their gene expression dynamics to variable energy budgets, this could theoretically occur in an unregulated manner simply due to ATP and metabolite availability. However, there might also exist regulatory mechanisms within GRNs that provide coupling of expression dynamics to energy budgets. One such mechanism has been described for developmental GRNs in *Drosophila* [9]. While using ablation of Dilp-secreting cells to reduce energy expenditure in larval cells, it was found that repressors of gene expression became dispensable for their regulatory functions on target genes. This phenomenon was so pervasive that when energy metabolism was reduced, the entire family of microRNA repressors could be eliminated with minimal effect on *Drosophila* development [9]. In contrast, under normal metabolic conditions, *Drosophila* microRNAs are essential for life [10].

A stylized mathematical model was developed to explain this phenomenon, predicated on the observed expression dynamics of many genes involved in development [9]. Developmental genes are often expressed in a succession of pulses, acting to successively restrict cell potential [11–14]. Each gene requires activators to induce an expression pulse and repressors to relax the pulse back to an off state. When energy metabolism is reduced, the kinetics of pulse relaxation are naturally slowed [4], mitigating the need for a full complement of repressors since repressor molecules have more time to completely act on their targets [9]. Remarkably, this stylized model appears to capture the dynamics due to changes in repressors of transcription, RNA processing/stability, and protein processing/stability. If the model, despite its simplicity, captures the processes at play, then it could explain why many genes are individually regulated by multiple repressors. When energy metabolism is elevated, auxiliary repressors would provide supplementary aid in relaxing expression pulses, thus giving cells greater robustness to fluctuations in nutrient availability.

In this study, we explore the possibility that activators of gene expression likewise become dispensable in developmental GRNs when energy metabolism is reduced. Both theoretical modeling and experiments indicate that this is indeed the case. We further provide evidence that accelerated metabolism results in a greater need for auxiliary gene activation. Thus, auxiliary gene activators provide greater robustness to fluctuations in nutrient availability.

## RESULTS

### A dynamical model describes the relationship between gene activation and energy metabolism

We used the previously developed mathematical model [9] to test the hypothesis that activators of gene expression become dispensable when energy metabolism is reduced. Conceptually, the model describes a simplified pathway of gene expression that represents the expression of a single gene within a succession of multi-gene pulses (Figure 1A,B). If the genes encode activators and repressors of other genes in the cascade, then simple GRN circuits can generate the pulsatile dynamics (Figure 1B). Indeed, such circuits are commonly found in developmental GRNs [11, 12, 14].

**Figure 1.**
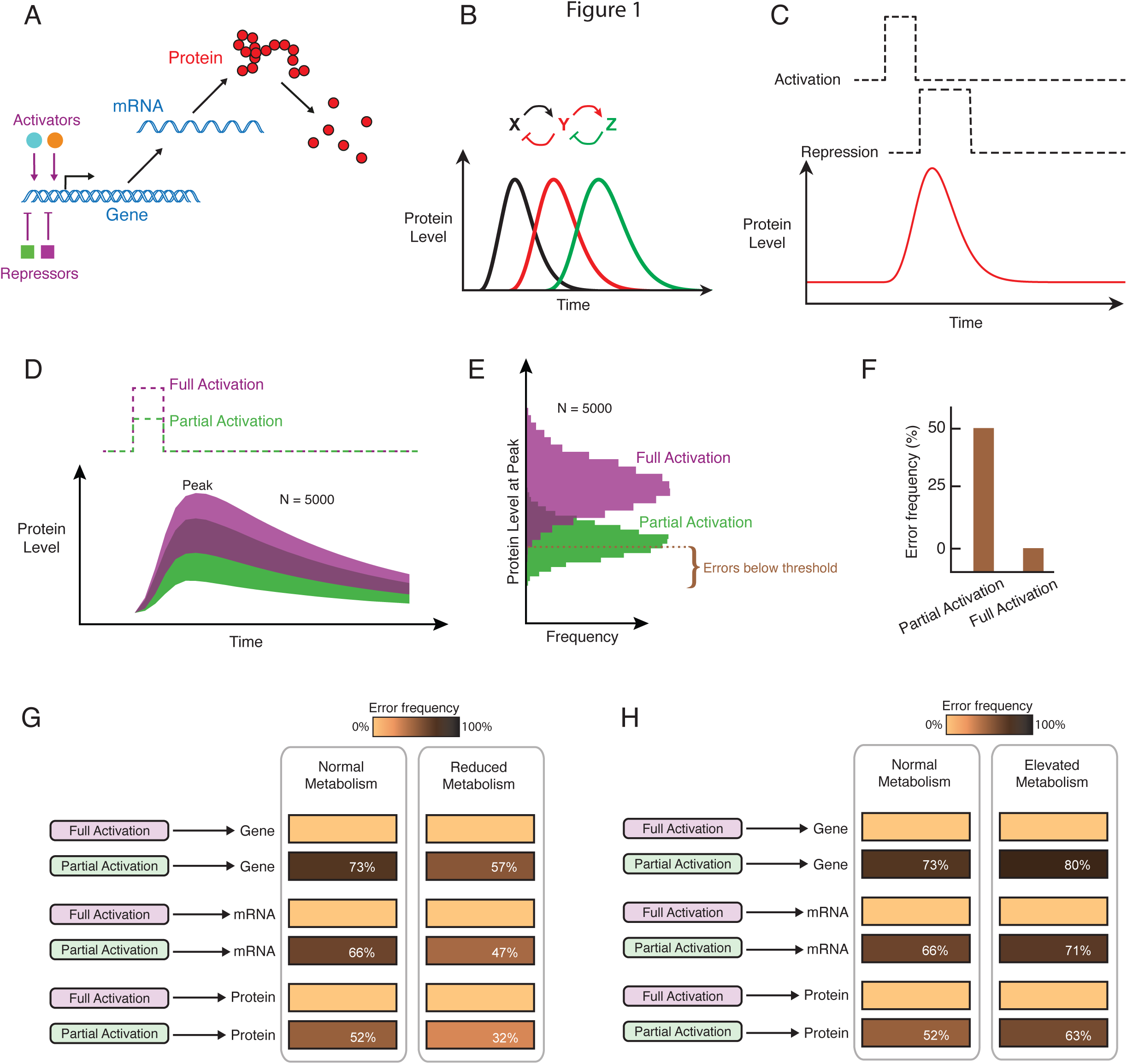
Mathematical modeling of gene activation and the effects of energy metabolism. **(A)** Schematic of generic regulation of gene expression. **(B)** Cascade of successive pulses in expression of three genes, whose products regulate each other as indicated at top. This program of gene expression occurs as a cell passes through a series of developmental states. The model focuses on transient expression of a single gene within the cascade. **(C)** Schematic of protein output from a single gene over time with a step function in gene activation followed by gene repression to relax expression to a baseline state. **(D)** Model simulations showing protein output over time. Shown are 5,000 simulated trajectories, which merge into a continuous band of trajectories. Green and purple denote simulations with 50% (partial) and 100% (full) activation of gene expression, respectively. **(E)** Frequency distribution of peak-level protein output from all simulations. A threshold is set where 99% of simulations with full activation are greater in peak output. **(F)** With partial activation, fewer simulations cross the threshold. Each failure to cross the threshold is an error. **(G)** Error frequency is greater with impaired activation at the transcriptional, RNA, and translational steps of gene expression. However, impaired activation imparts fewer errors when ATP-dependent parameter values are reduced by 50%. **(H**) Impaired activation imparts more errors when ATP-dependent parameter values are increased by 50%, regardless of how activators act on gene expression.

In our control theoretic model, activators transiently stimulate expression of a gene, whose protein is the output (Figure 1C). Each step of gene expression is potentially mediated by one or more activators acting in parallel: mRNA transcription, mRNA processing/stability, and protein translation. Combined, these activators help determine the size of the output’s pulse amplitude. Feedback repressors act to reduce the protein output, leading to a relaxation in output back to a baseline level (Figure 1C). Since gene expression is noisy [15], we used a stochastic simulation approach to infer the entire distribution of possible dynamic trajectories in protein output. There was a broad distribution of pulsatile trajectories from 5,000 such stochastic simulations (Figure 1D, purple traces).

We then compared the output dynamics when gene activation was reduced by 50% (i.e. one or more activators were absent) (Figure 1D, green traces). The two distributions partly overlapped but a large fraction of simulations with partial activation gave diminished protein output. This effect was consistently observed over a broad range of model parameter values, with 1,000 parameter sets tested (Figure S1A).

Since each pulse must trigger subsequent events in the GRN cascade, propagation of the cascade is contingent upon sufficient peak expression of each gene. Activators are critically important for the peak level (amplitude) of each pulse; peak output level was generally reduced when gene activation was reduced (Figure 1E). We defined a minimum amplitude that the output level must reach before a subsequent event is triggered. This threshold was defined such that exactly 99% of simulations with full activation achieved sufficient pulse amplitude to trigger a subsequent pulse (Figure 1E). The remaining 1% of trajectories that failed to reach the threshold level were denoted “errors”. Errors became much more frequent with partial gene activation (Figure 1E,F). This property was observed over a broad range of model parameter values (Figure S1B,C), and regardless of whether activators function in transcription, RNA processing, or protein translation (Figure 1G).

We next asked whether gene activation is less essential for peak protein output when energy metabolism is reduced. We reduced the rate parameters of each ATP-utilizing reaction by 50% to reflect conditions of reduced energy metabolism and compared simulation outcomes with full gene activation versus partial gene activation. The error frequency induced by partial gene activation was significantly diminished when ATP-dependent rate parameters were lowered (Figure 1G). This effect on activator loss occurred when activators function in transcription, RNA processing, or protein translation (Figure 1G), and it persisted across a wide range of parameter values (Figure 2A). The same effect was also evident when comparing cumulative output protein expression rather than the instantaneous peak levels reached by each expression pulse (Figure 2B). Furthermore, the effect persisted when (i) an upper bound of two alleles was placed on the gene’s transcription output (Figure 2C,D), (ii) cooperative transcription kinetics were taken into account (Figure 2E,F), and (iii) there was a nonzero basal level of gene expression (Figure 2G).

**Figure 2.**
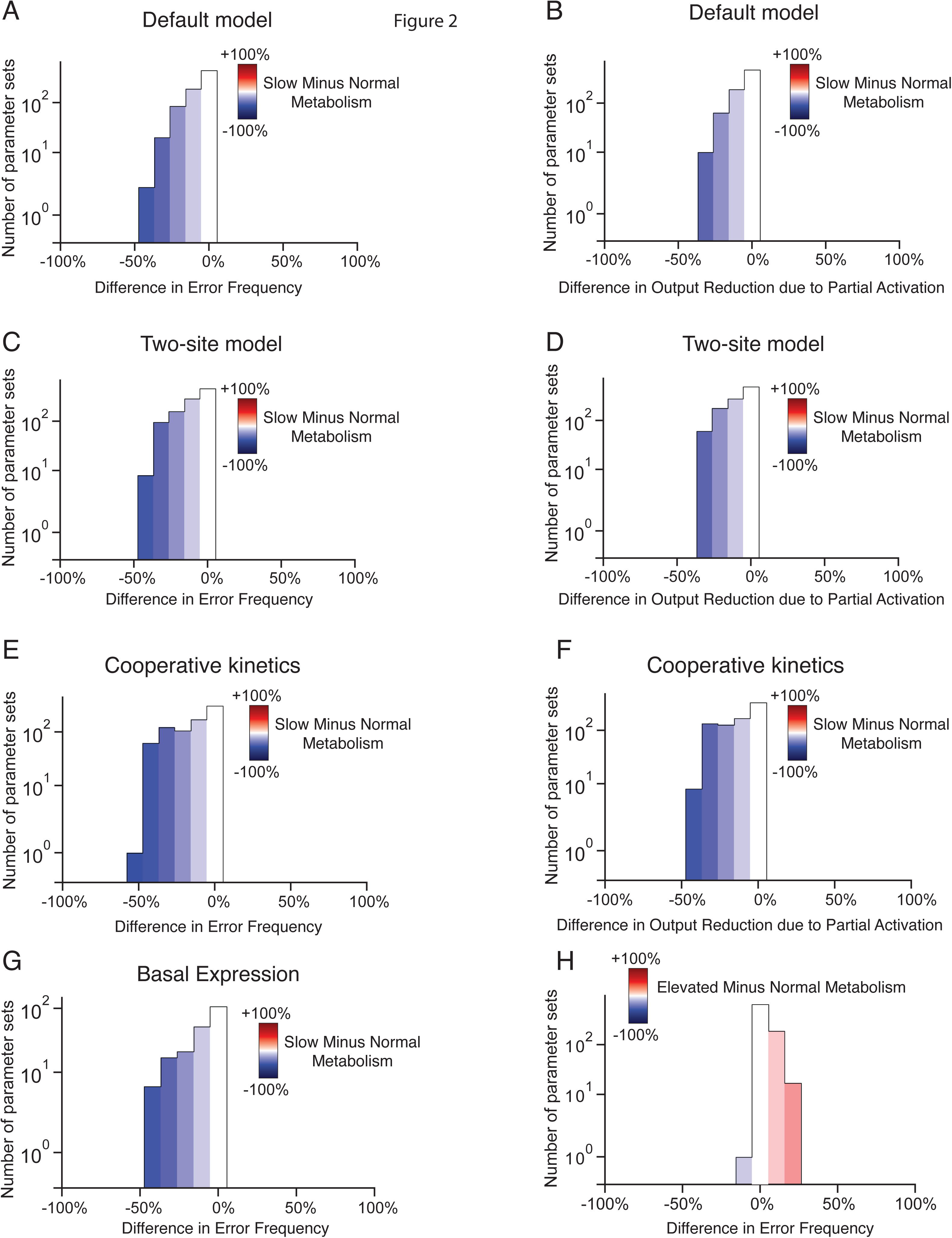
Model predictions are robust to model feature variation. Each of the seven model parameters was varied by one order of magnitude centered around the default value as defined in the STAR Methods. 1,000 parameter sets were generated, and 5,000 simulations with full and partial activation were performed for each parameter set. **(A-G)** Systematic modification of model conditions showing the difference between reduced versus normal metabolism for all parameter sets. For **(A,C,E,G)**, protein output at the peak of expression was compared between full and partial activation. Error frequency from partial activation was estimated using a threshold in peak expression. Shown are the distributions of the difference in error frequency between reduced and normal metabolism for all parameter sets. For **(B,D,F)**, protein output was calculated over the entire time course of gene expression, and the frequency in which output reduction occurred with partial activation was estimated for all parameter sets. Shown are the distributions of the difference in output reduction between reduced and normal metabolism for all parameter sets. **(A,B)** The default model. **(C,D)** Model where an upper bound of two is placed on the number of alleles transcribing the gene. **(E,F)** Model where cooperative transcription kinetics are considered. **(G)** Model where a nonzero basal stimulus is applied. **(H)** In the default model, error frequency from partial activation was estimated using a threshold in peak expression. Shown are the distributions of the difference in error frequency between elevated and normal metabolism for all parameter sets.

We then asked if elevating energy metabolism above normal exacerbates error frequency. To simulate the effect of elevating energy metabolism above normal, we increased the rate parameters of each ATP-dependent reaction by 50%, and found that error frequency was enhanced when gene activation was impaired (Figure 1H). This effect on activator loss occurred when activators function in transcription, RNA processing, or protein translation (Figure 1H), and it was observed across a wide range of parameter values (Figure 2H). Combined, all of our simulations predict that protein output of gene expression is differentially sensitive to changes in gene activation when energy metabolism is varied.

### Experimental validation of the dynamical model

We experimentally tested the model’s key prediction by measuring the expression dynamics of the regulatory protein Yan in *Drosophila*. Yan exhibits pulsatile expression in the *Drosophila* larval eye, where it is induced by a wave of cell-cell signaling that traverses the eye epithelium [16, 17]. Eye cells rapidly upregulate Yan protein abundance to a peak; the protein then decays back to initial levels over the course of 40 hours (Figure 3A). As cells achieve high levels of Yan, some of them are induced to transition to photoreceptor fates [17, 18]. However, Yan protein itself does not promote the transition but actually inhibits it through its repressive transcriptional activities [19].

**Figure 3.**
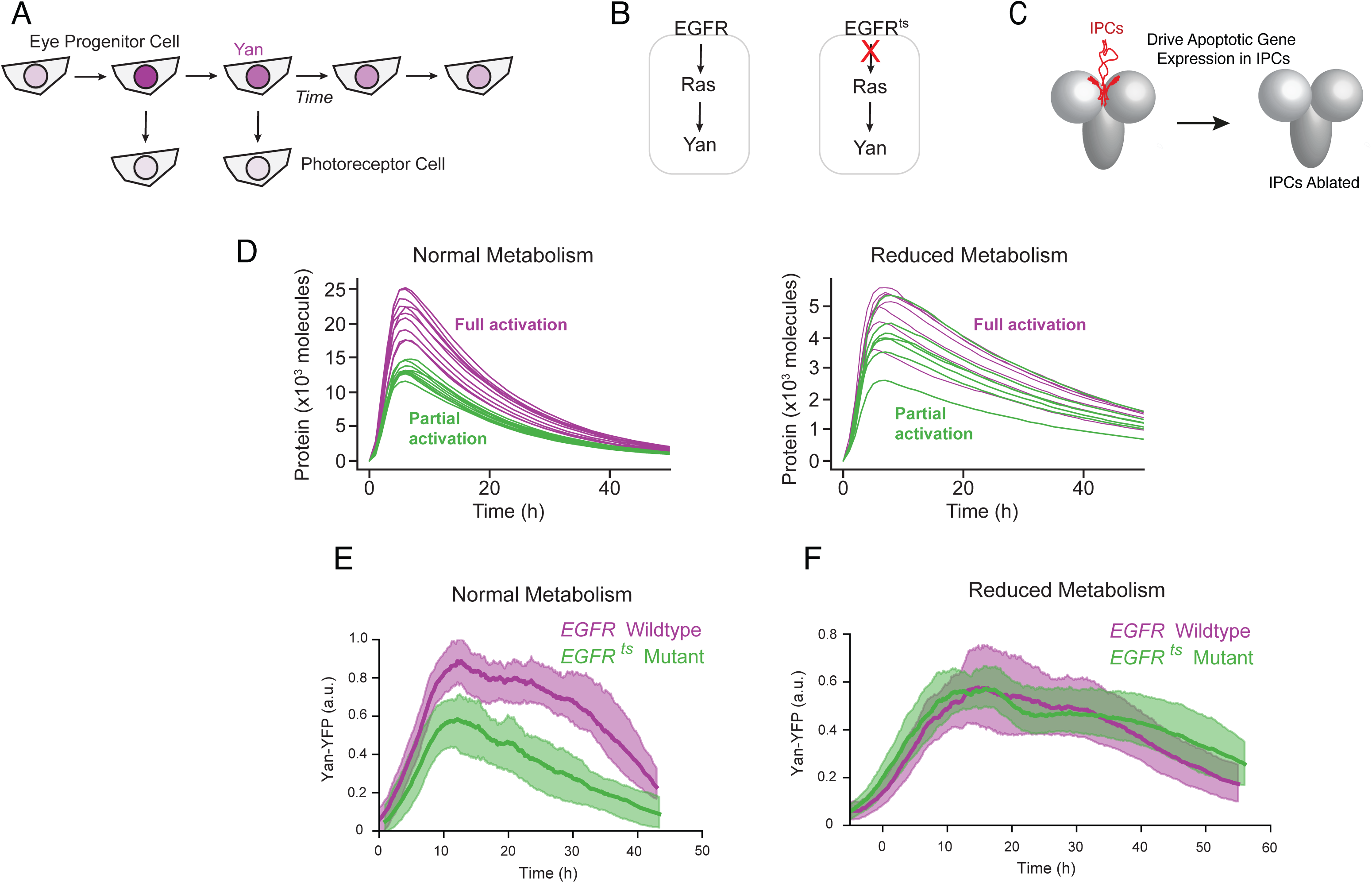
EGFR activation of Yan expression is dispensable when metabolism is slowed. **(A)** Schematic of Yan expression dynamics in eye disc cells. **(B)** Yan expression is positively dependent on EGFR in eye cells. **(C)** The 14 IPC cells in the larval brain are killed by specific expression of the pro-apoptotic protein Rpr. **(D)** Simulated protein output under the control of an auxiliary transcriptional activator (purple), and when the activator is removed (green). All simulations (purple and green) are also under control of a constitutive activator. Shown are ten randomly-chosen samples from a total population of 5,000 trajectories for each condition. Left, simulations performed with normal ATP-dependent reaction rates. Right, simulations performed following a 50% reduction in the rate of ATP-dependent reactions. **(E,F)** Yan-YFP expression dynamics in eye disc progenitor cells that are wildtype *EGFR* or ts-mutant *EGFR* raised at the non-permisssive temperature. Time 0 marks the time at which Yan expression begins. Solid lines are moving averages. Shaded regions denote 95% confidence intervals. Each line average is calculated from a composite of measurements of between 4,448 and 5,406 cells. **(E)** Yan-YFP dynamics under normal metabolic conditions. **(F)** Yan-YFP dynamics when the IPCs have been ablated.

To precisely measure Yan protein in cells, we used a *Drosophila* strain in which the protein is tagged with YFP and is still fully functional [16, 17]. Confocal microscopy of the eye discs was coupled with a computational pipeline for segmentation and analysis, yielding a composite picture of Yan dynamics sampled from thousands of cells per condition [17, 18].

The pulse of Yan expression in the eye is activated by signaling through the Epidermal Growth Factor Receptor (EGFR) (Figure 3B). When a temperature-sensitive (ts) *EGFR* mutant was transiently raised at a temperature of 26.5°C, peak output of Yan protein was reduced by ∼ 35% when compared to output at the permissive temperature of 18°C [17]. We repeated the experiment and observed a similar response in Yan output to the *EGFR* ts mutant (Figures S2 and S3). Thus, loss of EGFR activity reduces the peak level of Yan expression in the eye.

We then genetically ablated the insulin-producing cells (IPCs) of the larval brain (Figure 3C), which causes a 40% reduction in cell uptake of circulating sugars, a decrease in mitochondrial oxidative phosphorylation, and a 70% slowdown in overall fly development [9, 20]. Model simulations predicted that the amplitude of protein output will be less sensitive to impaired gene activation if energy metabolism is reduced (Figure 3D). We ablated the IPCs and observed that loss of EGFR had little to no impact on Yan expression when the IPCs were ablated (Figure 3E,F). This behavior clearly resembled the simulated dynamics under conditions of reduced energy metabolism (Fig. 3D).

The model also predicted that elevating metabolism to above-normal levels would enhance the dependence of gene expression on activators (Figures 1H and 2H). To test this prediction, we overexpressed the Myc transcription factor in eye cells. This induced elevated cellular growth and reconfigured cellular metabolism so that oxidative phosphorylation is displaced by aerobic glycolysis [21–24]. Myc overexpression also increased mitochondria number in cells [24], which we observed when overexpressing Myc in *Drosophila* cells (Figure 4A). We specifically overexpressed Myc in eye epithelial cells that were *EGFR* mutant, and we observed a greater impact of EGFR on Yan output. While peak expression was reduced by 34% in a wildtype Myc background, (Figure 4B), peak expression was reduced by 45% in the Myc overexpression background (Figure 4C). Thus, modulating metabolic activity to be lower or higher than normal demonstrates how activators have variable effects on target gene expression depending on metabolic conditions.

**Figure 4.**
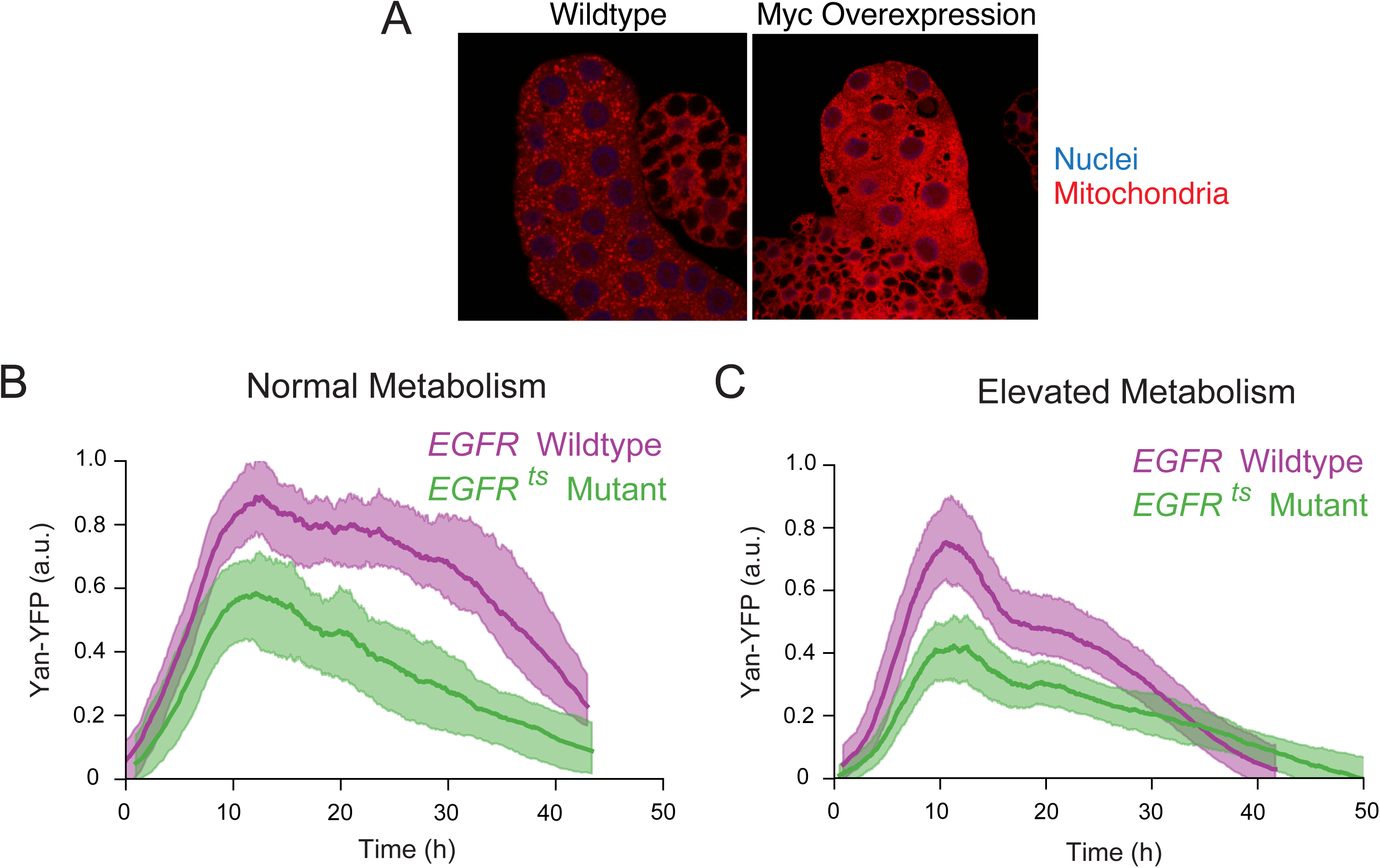
Yan expression is more dependent on EGFR when metabolism is elevated. **(A)** Larval salivary gland cells from *ptc-Gal4/+* (left) and *ptc-Gal4/UAS-Myc* (right) animals. Tissue was stained for mitochondria (red) and nuclei (blue). Note the higher density of mitochondria in cells overexpressing Myc. **(B,C)** Yan-YFP expression dynamics in eye disc progenitor cells that are wildtype *EGFR* or ts-mutant *EGFR* raised at the non-permisssive temperature. Time 0 marks the time at which Yan expression begins. Purple and green solid lines are moving averages of wildtype and mutant cells, respectively. Shaded regions denote 95% confidence intervals. Each line average is calculated from a composite of measurements of between 4,448 and 8,653 cells. **(B)** Yan-YFP dynamics under normal metabolic conditions. **(C)** Yan-YFP dynamics in *GMR-Gal4/UAS-Myc* eye discs. Solid lines are moving averages. Shaded regions denote 95% confidence intervals. Although it appears that overall Yan-YFP output is not increased with Myc overexpression, it is important to keep in mind that Yan repressors are also likely more active, contributing to a more normalized expression output.

### Developmental outcomes are dependent on metabolism - gene activator interactions

We had previously found that ablation of IPCs suppressed developmental phenotypes caused by mutations in gene repressors [9]. Therefore, we wondered if IPC ablation also suppressed phenotypes caused by loss of activators. Yan is important for the proper specification of photoreceptors in the eye [19], and transient loss of EGFR activity results in mispatterned adult compound eyes in which the highly regular hexagonal lattice of unit eyes (ommatidia) becomes disordered [17, 25]. To precisely measure the degree of disorder in such eyes, we developed a new image-based analysis pipeline. Brightfield microscopy of adult compound eyes captured the reflection points of individual ommatidium lenses (Figure S4A-E). After computational segmentation of reflection points, triangulation of their centroids allowed us to measure the distance between each ommatidium and all its immediately adjacent neighbors (Figures 5A and S4F-I). For each ommatidium, we calculated the difference between its distance to its closest neighbor and its distance to its farthest neighbor. This value *D*, normalized by the average interommatidial distance, was calculated for all ommatidia in all eye samples (Figure 5B). The *D* metric is a measure of lattice disorder. Errorless measurements on a perfectly regular lattice yield an average *D* of zero; measurement errors on the order of 10% of the average distance yield a *D* ≈ 0.15. Disordered lattices will yield even higher values for *D* (Figure 5C). We applied the method to measure disorder in the compound eyes of wildtype and mutant flies. Both EGFR and the miRNA miR-7 regulate expression of Yan in the eye [17, 26]. Visual inspection of *EGFR* and *mir-7* mutant eyes showed a qualitatively greater disorder compared to wildtype (Figure S5). Quantitatively, both the *EGFR* and *mir-7* mutant adults exhibited greater eye lattice disorder since their mean values for the *D* metric were significantly higher than wildtype controls (Figure 5D,E).

**Figure 5.**
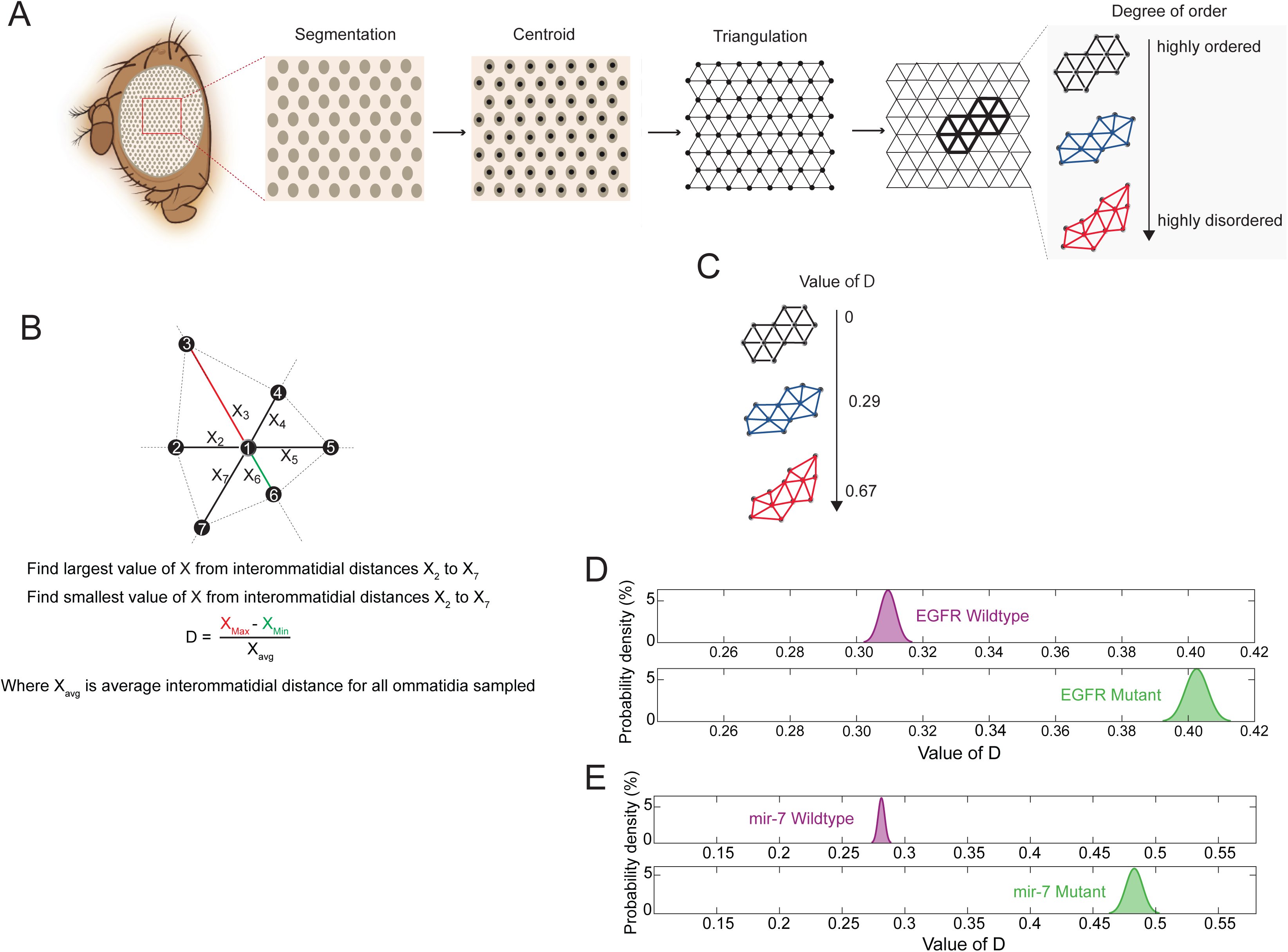
Quantitation of lattice disorder in mutant compound eyes. **(A)** Pipeline of analysis involves segmentation of ommatidia, triangulation of their centroids to recreate the overall lattice, and local lattice analysis of every ommatidium and its nearest neighbors. **(B)** Local lattice disorder is estimated as the difference between the longest distance from one ommatidium to a nearest neighbor and the shortest distance from that ommatidium to a nearest neighbor. This normalized difference *D* is calculated for every ommatidium in the region of interest. **(C)** Schematic of ommatidia with varying values of *D* and therefore varying levels of disorder. **(D)** Density distributions of the mean *D* estimated for ommatidia from wildtype (purple) and *EGFR ts* mutant (green) eyes. Animals were raised at 18°C except for an 18-hour interval as late L3 larvae when they were incubated at 26.5°C. The numbers of ommatidia analyzed for each dataset were 1,321 and 1,438, respectively, and these were each imaged from 10 animals. **(E)** Density distributions of the mean *D* estimated for ommatidia from wildtype (purple) and *mir-7* mutant (green) eyes. The numbers of ommatidia analyzed for each dataset were 1,333 and 1,130, respectively, and these were each imaged from 10 animals.

We then measured eye lattice disorder in *EGFR* mutants that had their IPCs ablated (Figures 6A and S6). The mean *D* metric was significantly lower in *EGFR* mutants with IPC ablation than in *EGFR* mutants with normal metabolism. Hence, lowering energy metabolism suppresses the developmental phenotype of the *EGFR* mutant. We also measured disorder in *EGFR* mutants that overexpressed Myc in eye cells (Figures 6B and S6). There was a significant increase in the mean *D* metric relative to controls with normal metabolism. Thus, raising metabolism enhances the *EGFR* developmental phenotype. Overall, these results are consistent with the results of Yan expression experiments, implicating energy metabolism as a modulator of gene regulation through activation.

**Figure 6.**
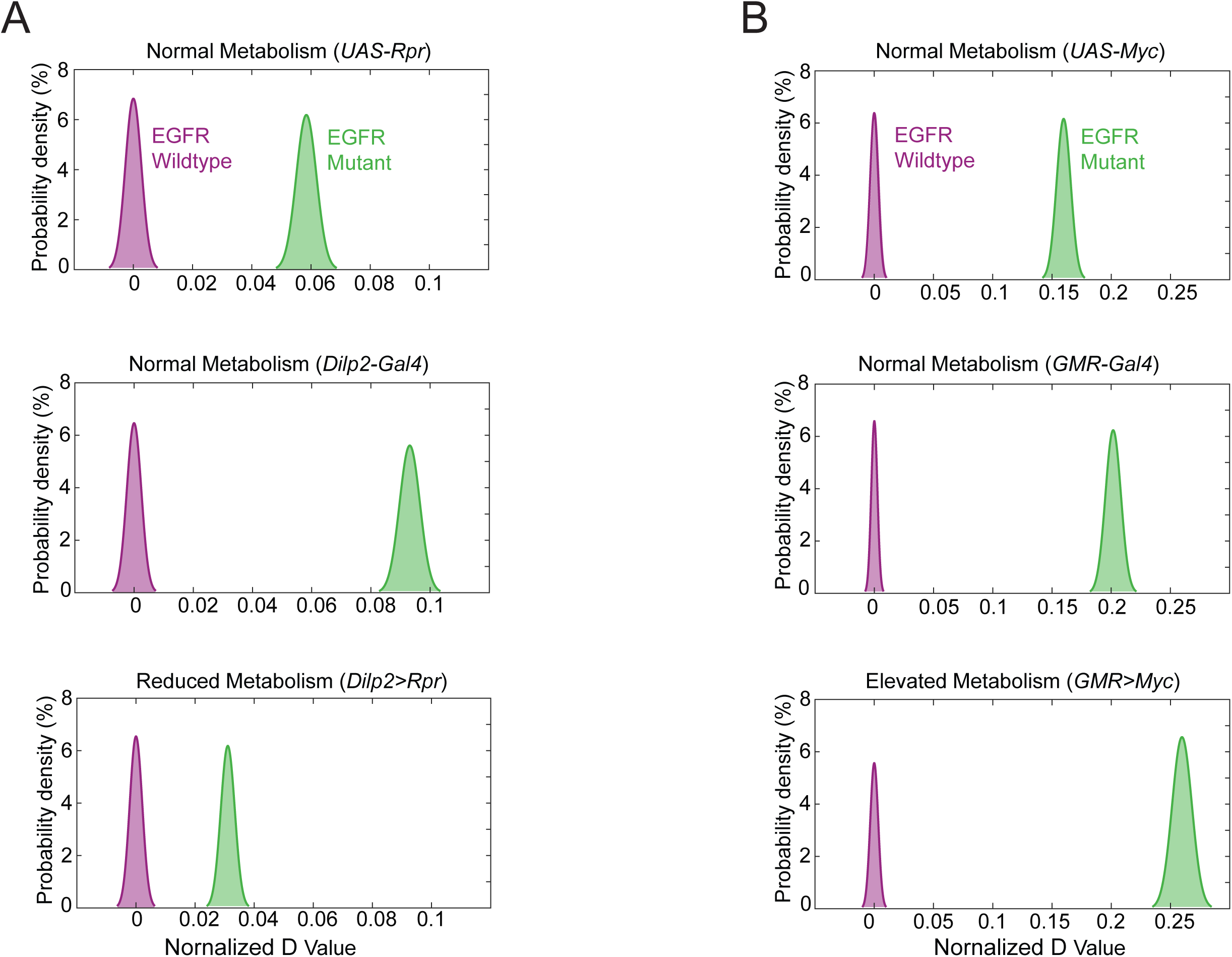
EGFR loss affects disorder of the eye lattice dependent on energy metabolism. **(A,B)** To compare between *EGFR* genotypes, the density distribution of the mean *D* estimated for ommatidia from wildtype *EGFR* eyes (purple) was adjusted to center around zero, and the *EGFR* mutant distribution (green) was normalized accordingly. **(A)** Disorder of *EGFR* mutants when metabolism is normal or is reduced by IPC ablation. **(B)** Disorder of *EGFR* mutants when metabolism is normal or is elevated by Myc overexpression.

## DISCUSSION

Development and growth are fueled by energy metabolism suggesting that the tempo of development depends on metabolic rate. Thus, the dynamics of developmental GRNs must faithfully adjust to a variable time scale. We have shown that auxiliary activation of gene expression within an eye developmental GRN synchronizes expression dynamics with the variable pace of the developmental program. Activation by EGFR is required for proper Yan expression and developmental outcome when energy metabolism is normal, and EGFR becomes functionally redundant when metabolism is lowered. A stylized mathematical model predicts that this relationship between gene activators and metabolism is not limited to Yan expression but exists for many genes and their activation factors. In this way, auxiliary gene activation would allow for reliable development across a broader range of metabolic conditions than would otherwise be tolerated.

A similar relationship between metabolism and gene repressors has also been documented [9]. The intuition for understanding this relationship resides in the fact that limiting ATP availability reduces the rates of anabolic reactions but not all catabolic reactions. This asymmetric effect on different steps in gene expression is one reason why gene repression becomes less essential when ATP availability is limited. Downregulation of protein levels are more dominated by constitutive decay processes when ATP becomes limiting. Another reason repression becomes less essential is that the dynamics of molecular synthesis and degradation become slower, enabling the overall control mechanism to adjust protein levels with less negative feedback. The reason why activation becomes less essential with limiting ATP is because the slower dynamics of synthesis and degradation also allow for weaker activation to nevertheless achieve peak output despite negative feedback control. Our modeling predicts that reducing metabolism does not rescue loss of activators as extensively as it rescues loss of repressors. Indeed, the eye phenotype when EGFR is lost is only partially rescued by IPC ablation whereas the eye phenotypes of repressor mutants are strongly rescued by the same ablation [9].

Our model is not limited to developmental genes but could be relevant for any genes that are expressed with pulsatile dynamics due to negative feedback control. Many genes in both unicellular and multicellular organisms have these features [27, 28]. Examples include stress response genes, and genes involved in signal transduction [29, 30]. It will be interesting to discover whether our model predictions of metabolism and gene control extend to such situations.

The insect eye is an example of a system with remarkable spatial order that is driven by a demand for optimal visual acuity and sensitivity [31, 32]. We have developed an imaging-based analysis tool that precisely measures spatial order in the *Drosophila* compound eye. The method is readily adaptable for general use and is potentially applicable for insect species other than *Drosophila melanogaster*. Here, we apply it to measure disorder in genetic mutants, but it can be used to study the effects of other perturbations, and its sensitivity will potentially be useful for even the weakest of perturbations.

## ACKNOWLEDGEMENTS

We thank the NIH (GM118144, R.W.C.), the NSF (1764421, L.A.N.A., N.B. and R.W.C), and Simons Foundation (597491 L.A.N.A., N.B. and R.W.C) for funding, Bob Holmgren and the Bloomington Drosophila Stock Center for fly stocks, Laura Nilson for the use of computational resources, and the Northwestern Biological Imaging Facility (BIF) for technical imaging support.

## AUTHOR CONTRIBUTIONS

The experimental work was conceived by S.Q. and R.W.C. All eye experiments were performed by S.Q., under the supervision of R.W.C. K.D.G. designed and built the computational pipeline for segmentation of compound eye ommatidia and the quantification of lattice disorder, with assistance from S.Q, and under the guidance of R.W.C. S.Y. performed all imaging and analysis of mitochondria in cells overexpressing Myc. The modeling was conceived by S.B., N.B., and L.A.N.A. All model analysis was performed by S.B. under the supervision of N.B. and L.A.N.A. The manuscript was written by R.W.C. with input from all authors.

## METHODS

### Drosophila Growth and Genetics

For all experiments, *Drosophila melanogaster* was raised using standard lab conditions and food. All experiments used female animals unless stated otherwise. Stocks were either obtained from the Bloomington Stock Center, from listed labs, or were derived in our laboratory (RWC). Experiments with *EGFR* were performed using trans-heterozygous mutants in order to minimize phenotypes induced by secondary mutations on relevant chromosomes. Trans-heterozygous allele combinations used were *Egfr^tsla^*/*Egfr^f24^* [17]. Genetic mosaic animals bearing *mir-7*^△^*^1^* homozygous mutant eyes were generated using the ey-FLP/FRT system as described [9].

The BAC genomic transgene *Yan-YFP* was inserted on chromosome 3 at the attP2 site. The *Yan-YFP* chromosome was homozygosed so that animals had two copies of the transgene, and placed in a *yan^443^ / yan^884^* mutant background so that the endogenous *yan* gene did not make any protein.

To genetically ablate the insulin producing cells (IPCs) of the brain, *yw* animals were constructed bearing an *Dilp2-Gal4* transgene on chromosome III and a *UAS-Reaper* (*Rpr*) transgene on chromosome I. *Rpr* is a pro-apoptotic gene that is sufficient to kill cells in which it is expressed [33]. *Dilp2-Gal4* fuses the *insulin-like peptide 2* gene promoter to Gal4, and specifically drives its expression in brain IPCs [20]. Examination of *Dilp2-Gal4; UAS-Rpr* larval brains showed that they almost completely lacked IPCs [9]. Previous studies found that IPC-deficient adults are normally proportioned but of smaller size. It takes 70% longer to complete juvenile development, and juveniles have a 40% elevation in blood glucose, consistent with insulin-like peptides (Dilps) being essential regulators of glucose metabolism in *Drosophila* [9, 20, 34, 35]. Moreover, animals generate 30% less heat output as measured by whole-body calorimetry [8], and ATP synthase abundance is reduced in cells from IPC-ablated larvae [9]. For all wildtype controls, we tested animals bearing either the *Dilp2-Gal4* or *UAS-Rpr* gene in their genomes.

To overexpress the *Drosophila* transcription factor Myc in cells, animals were constructed with a *UAS-Myc.Z* transgene located on chromosome II or III. This transgene was activated using either a *GMR-Gal4* transgene or *ptc-Gal4* transgene located on chromosome III or II. *GMR-Gal4* drives UAS gene expression in all larval eye cells posterior to the morphogenetic furrow [36], while *ptc-Gal4* drives UAS gene expression in larval salivary gland cells [37]. Ptc>Myc causes salivary gland cells to grow faster due to enhanced translation capacity [38]. GMR>Myc causes eye cells to grow 33% bigger [23]. Myc overexpression reconfigures cellular metabolism in imaginal discs so that oxidative phosphorylation is displaced as the predominant source of energy production by increased glycolysis, resembling the Warburg effect [24]. Expression of Myc increased the mitochondrial network, consistent with Myc’s regulation of mitochondrial biogenesis [24].

EGFR activity was conditionally reduced by placing *Egfr^f24^* [39] *in trans* to the ts mutant allele *Egfr^tsla^* [25]. Flies were raised at the permissive temperature (18°C) and shifted to a semi-restrictive temperature (26.5°C) as third instar larvae for 18 hours. Genetic wildtype controls were heterozygotes of either *Egfr^f24^* or *Egfr^tsla^* over a wildtype chromosome. These controls were also shifted to 26.5°C as third instar larvae for 18 hours. At 26.5°C, developing eye cells had compromised EGFR activity in *Egfr^tsla^*/*Egfr^f24^* animals since when transferred back to the permissive temperature and allowed to eclose, they had rough eyes [17]. In mutant eye discs, there were signs of some cells undergoing apoptosis - a significant reduction of nuclear diameter, a strong Yan-YFP brightness, and anomalous nuclear position along the apical-basal axis. Apoptosis was more prevalent in discs from mutant animals treated at temperatures greater than 26.5°C [17]. Therefore, we chose this temperature for EGFR activity reduction so as to minimize apoptosis but still achieve effects on Yan expression. We only included in our analysis cells corresponding to classical anatomical positions and apical basal migration patterns.

### Quantification of Yan-YFP Expression Dynamics in the Eye

Eye discs from white-prepupae of the correct genotype (*yan^443^ / yan^884^; Yan-YFP / Yan-YFP*) were dissected, fixed, and imaged by confocal microscopy, as previously described [17, 18]. These prepupae also had the appropriate combinations of *Egfr* alleles, *Gal4* driver genes, and *UAS* transgenes for controlled manipulation of EGFR activity and cell metabolism. Eye discs were fixed for 45 min at room temperature in 4% (w/v) paraformaldehyde in PBS. Discs were washed in PBS and then incubated in 1:1 (v/v) PBS:VectaShield with DAPI (Vector Laboratories) for 45 min, followed by a 45 min incubation in 100% VectaShield with DAPI. Samples were then mounted with VectaShield with DAPI and imaged by Leica TCS SP8 confocal microscopy equipped with a 40X oil objective (NA = 1.3) with a digital zoom of 1.2 - 1.4. Yan-YFP and DAPI were separately detected by HyD detectors (GaAsP). During imaging, discs were oriented with the morphogenetic furrow parallel to the y axis of the image. Optical slices were captured as 1024 × 1024 8-bit images, in which at least 6 rows of ommatidia on either side of the dorsal-ventral equator were recorded. Optical slices were set at 0.7 μm thickness, and 45 - 60 optical slices were captured in a z-stack to completely image eye discs from basal to apical surfaces. All discs for a given condition were fixed, mounted, and imaged in parallel to reduce measurement error. Sample preparation, imaging, and analysis were not performed under blind conditions. See Figure S3 for examples of typical imaging data.

Image data was processed for automatic segmentation and quantitation of DAPI and YFP nuclear fluorescence as described [17, 18]. Briefly, cell segmentation was performed using the DAPI signal as a reference channel for identification of cell nuclei boundaries. Each layer of the reference channel was segmented independently. A single contour containing each unique cell was manually selected and assigned a cell type using a custom graphic user interface called *Silhouette*. For each annotated cell contour, expression measurements were obtained by normalizing the mean pixel fluorescence of the YFP channel by the mean fluorescence of the DAPI channel. This normalization serves to mitigate variability due to potentially uneven sample illumination, segment area, and differences in protein expression capacity between cells. We assigned cell-type identities to segmented nuclei by using nuclear position and morphology, two key features that enable one to unambiguously identify eye cell types without the need for cell-specific markers [40]. This task was accomplished using *Silhouette*; an open-source package for macOS that integrates our image segmentation algorithm with a GUI for cell type annotation. Subsequent analysis and visualization procedures were implemented in Python using the *FlyEye* package, open source software developed by our group.

Using *FlyEye*, cell positions along the anterior-posterior axis were mapped to developmental time as described previously [17, 18]. This is predicated on two assumptions: the furrow proceeds at a constant rate of one column of R8 neurons per two hours; and minimal cell migration occurs. For each disc, Delaunay triangulations were used to estimate the median distance between adjacent columns of R8 neurons [41]. We used the median rather than the mean distance because it minimized the influence of non-adjacent R8s that were falsely identified by the triangulation [17]. Dividing the furrow velocity of 2 h per column by this median distance yields a single conversion factor from position along the anterior-posterior axis to developmental time. This factor was applied to all cell measurements within the corresponding disc, yielding expression time series. Notably, these are not single cell dynamics, but rather aggregate dynamics across the development time course of a spatially organized cell population.

Moving averages were computed by first-order Savitzky-Golay filtration [42]. This method augments the simple windowing approach used in [17] by enabling visualization of expression trends at early time-points that are otherwise obscured by large window sizes. A secondary first-order filtration with one-fifth the original window size was applied to smooth lines for visualization purposes. None of our conclusions are sensitive to our choice of filtration or smoothing method. A primary window size of 250 cells was used for reporting the expression of cells, unless noted otherwise. Confidence intervals for the moving average were inferred from the 2.5th and 97.5th percentile of 1,000-point estimates of the mean within each window. Point estimates were generated by bootstrap resampling with replacement of the expression levels within each window.

To align multiple eye disc samples using *FlyEye*, cells of each sample were aligned with a reference population by shifting them in time as described in Bernasek et al [18]. The magnitude of this shift was determined by maximizing the cross-correlation of progenitor Yan-YFP expression *Y(t)* with the corresponding reference time series *X(t)*. Rather than raw measurements, moving averages within a window of ten cells were used to improve robustness against noise. This operation amounts to:

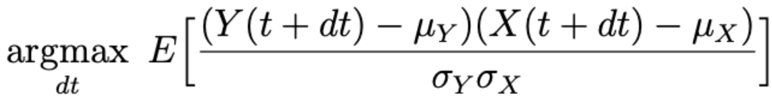

where, *μ* and *σ* are the mean and standard deviation of each time series, and *dt* is the time shift by which the population should be shifted.

For each experimental treatment, a disc was randomly chosen and shifted in time such that time “zero” corresponds to the first annotated R8 neuron. This disc then served as the reference population for the alignment of all subsequent biological replicates within the treatment. Similarly, different experimental treatments (e.g. control and perturbation) were aligned by first aligning the discs within each treatment, then aggregating all cells within each treatment and repeating the procedure with the first treatment serving as the reference. To plot the moving line averages of each aggregate dataset, we adjusted time on the x axis such that −10 hours became the new 0 hours and all other time intervals were adjusted accordingly. Yan-YFP is first detectable in cells no earlier than the −10 hour timepoint.

We analyzed six to twelve replicate eye discs for each treatment in two separate experiments. In total, we measured 5,406 and 4,870 cells under normal metabolism for *EGFR* wildtype and *EGFR* mutant samples, respectively. We measured 4,448 and 5,186 cells from *Dilp2>Rpr* animals for *EGFR* wildtype and *EGFR* mutant samples, respectively. We measured 4,668 and 8,853 cells from *GMR>Myc* animals for *EGFR* wildtype and *EGFR* mutant samples, respectively.

### Imaging of Adult Compound Eyes

For imaging compound eyes, 2-to 3-day-old adults of different genotypes were collected and stored in 100% ethanol. Before imaging, samples were progressively rehydrated by successive 24-hour incubations in 75% ethanol, 50% ethanol, 25% ethanol, and water. Blu-Tack (Bostik Smart Adhesives) was cut into a 1.5 cm piece and pressed using a thumb onto a microscope slide. The rehydrated flies were transferred onto a Kimwipe to briefly dry, and then were placed laterally onto the Blu-Tack with their left eyes facing up and oriented horizontally (Figure S4A). Mounted animals were imaged with a Leica DM6B bright field microscope with a 10x objective (NA = 0.40) and DFC7000T camera. To illuminate the samples, gooseneck fiber-optic lights (Schott, KL 1500 LCD) were positioned above the stage on opposing sides of the specimen. The two lights were positioned facing one another, and the angle of the fiber-optic cables was parallel to the stage (Figure S4B). Since image quality is affected by positioning of the light source, a criterion for a good quality image is determined by avoiding 1) uneven distribution of the light, 2) double reflective images from each ommatidium, and 3) reflection from non-ommatidia regions (Figure S4D,E). All objectives except the one in use must be removed from the microscope to prevent these lighting/imaging aberrations. In addition, the gooseneck lights must be subtly adjusted for each sample to avoid lighting aberrations. Exposure time was adjusted according to the eye color and eye size of each sample to ensure a uniform field of reflective points. An example of a high-quality image is shown in Figure S4C. Optical slices were captured at 10 μm intervals along the z-plane using the Leica Application Suite X. The stack of raw image files for each sample was imported into ZereneStacker (Zerene Systems, Richland, WA), from which DMAP images were generated as described [43]. Zerene Stacker projects the entire stack into a DMAP image. The DMAP files from ZereneStacker were then used for further analysis.

### Pipeline for Quantification of Eye Disorder

To computationally segment ommatidia, the raw DMAP images were pixel-classified into two classes of pixels: (1) pixels representing light reflected from ommatidia (identified by the reflection from each ommatidia lens), (2) pixels representing all other features of the images. This pixel classification step was achieved using the ‘Pixel Classification’ workflow in Ilastik, which is open-source software that provides machine learning image analysis [44]. Briefly, Ilastik was trained on 12 DMAP images from six different genotypes (two images were chosen from each genotype). This was to ensure good representation of the variability contained within the complete dataset. For the 12 images, pixels were manually annotated as belonging to the light reflected from ommatidia vs. not. This process was repeated until the model learned to satisfactorily classify pixels, determined using the live prediction feature of Ilastik. Once a satisfactory model was trained, the remaining 48 images in each dataset were pixel-classified using the trained Ilastik model.

Custom MATLAB scripts were developed that (1) import the pixel classification maps generated by the model trained in Ilastik, (2) threshold the pixel classification maps to obtain binary maps where 1’s represent the ommatidia light reflections, and 0’s represent all other pixels, (3) detect all isolated binary objects (contiguous pixels of value 1 that correspond to each ommatidium), (4) compute the centroid of each binary object, which then becomes the defined center of each ommatidium. Because each binary object was typically 10-20 pixels in size, it necessitated calculation of the object’s centroid to have a single pixel that defines the location of each ommatidium.

Since ommatidia were detected via light reflections, the automated workflow led to some misclassification of other reflections as originating from ommatidia. These reflections often occurred at ommatidial boundaries (particularly for rough eye phenotypes) and outside the eye field on the cuticle of the head. Therefore, a custom GUI was developed in MATLAB that allowed for manual correction of the data derived from the above described workflow. Briefly, this GUI displayed the DMAP images overlaid with all identified centroids. It allows the user to add new centroids for ommatidia that were not classified as ommatidia. It also allows the user to delete centroids that were improperly classified as ommatidia. This GUI was used to thoroughly correct the classification of ommatidia from the data.

For each analyzed image, the ommatidial lattice was defined using Delaunay triangulation of the centroids, implemented using a built-in MATLAB function. Because a triangular lattice is dual to a hexagonal lattice, Delaunay triangulation allows for analysis of the component triangles of each hexagonal unit of the compound eye – i.e. for wildtype eyes, each ommatidial unit is composed of six triangles that define the space between one ommatidia and its six closest neighbors; together, these six triangles create one hexagonal unit.

Interommatidial distances were calculated for all ommatidia except for those along the boundary of the region of analysis. This is because ommatidia along the boundary had neighbors that were not identified, preventing proper calculation of their local disorder. Boundary ommatidia did, however, contribute to the calculation of local disorder for ommatidia to which they were neighbors. Boundary ommatidia were identified by finding the convex hull of segmented ommatidia using the built-in MATLAB function.

Since we had generated a 2D image of a 3D curved structure, there were inherent distortions of the derived lattice that contributed to our estimation of lattice disorder. To minimize this curvature-based distortion of interommatidial distances, we only analyzed ommatidia within a fixed distance from the center of each eye, where there was minimal curvature. The distance chosen was 200 pixels and the center of each eye was estimated as the center of mass of segmented ommatidia. By limiting the analysis to a small region of the eye, effects of curvature were minimized.

Definition of the ommatidial lattice by Delaunay triangulation creates several easy-to-measure features of lattice order. These features include 1) the length of each triangle edge, 2) the area of each triangle, and 3) the angles formed at the vertices of each triangle. In a perfectly regular lattice, each of these three features would be uniform, i.e., all sides of every triangle would be the exact same length. We initially calculated lattice regularity by analyzing the distributions of the aforementioned lattice features. Variation was estimated as either the Coefficient of Variation or the Fano Factor. Although these estimates are descriptive of lattice order / disorder, they are not sensitive to infrequent or mild lattice defects.

Therefore, we devised a more sensitive measurement of lattice order. Local regularity of the lattice was defined as the variability in the distances that connect one ommatidium to its nearest neighbors. By finding the largest (*X_max_*) and smallest (*X_min_*) distance connecting one ommatidium to its nearest neighbors, the difference in those two distances is a measure of local order. For example, if *X_max_* - *X_min_* is zero, then the local lattice has perfect order. The larger the value for this difference, the greater the local disorder. We then normalized the difference to estimate

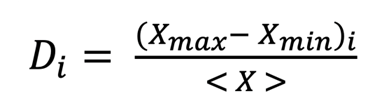

where *D_i_* is the degree of disorder for ommatidium *i*, and *<X>* is the mean distance between every ommatidium and its nearest neighbors. This *D* metric was calculated for every ommatidium analyzed in all eye samples from a given condition.

Another benefit of analyzing the regularity of the lattice on such a local scale is that it controls for distortion from eye curvature of the eye. When we calculated lattice order using the aforementioned three lattice features described above, a large contributor to the variability came from curvature-based distortion. Since *D_i_* is a local measurement, the scale of curvature is much greater than the scale of the measurement. Therefore, curvature contributes much less to the variability of *D_i_* measurements.

For statistical analysis, we performed bootstrapping on the thousands of measurements of *D* for each genetic condition (ranges from 993 to 1,473). Bootstrapping was performed in MATLAB for 10,000 times, and the mean value of *D* was calculated per bootstrap sample. The distribution of means was plotted as a histogram, and shown in the figures are the smoothed fits to each of the histograms.

The complete computational pipeline for segmentation, correction, and analysis that is described above is available as a MATLAB software package called *roughEye*, and is freely available.

### Mitochondria Staining

Second instar larvae (either *ptc-Gal4* genotype or *ptc-Gal4/UAS-Myc* genotype) were dissected, and salivary glands were incubated in PBS supplemented with 500 nM MitoTracker Red CMXRos (Thermo Fischer) for 30 min at room temperature. Glands were then washed several times with Schneiders medium for 10 minutes at room temperature. Glands were fixed in 4% (w/ v) paraformaldehyde in PBS for 20 minutes. After washing in PBS, glands were mounted in VectaShield (Vector Labs) with DAPI (to visualize nuclei). The samples was imaged with a Leica SP5 confocal microscopy system.

### Mathematical Modeling

Our modeling framework is based on the one we developed previously [9]. It directly describes the emergent expression dynamics of a single gene within a cascade of developmental gene expression. It leverages two key concepts from control theory. The first is the notion of Lyapunov stability; that is, systems tend to remain near stable equilibria. The second is the Hartman-Grobman theorem, which posits that systems deviate approximately linearly about these fixed points [45]. We therefore developed a model that describes the time evolution of linear deviations about the basal protein level that exists before gene expression is induced and after it subsides.

Specifically, a linear time invariant system describes the time evolution of deviations (*Δ)* in activated DNA (*ΔD*), mRNA (*ΔR*), and protein (*ΔP*) state variables in response to a change in stimulus (*ΔI*) that induces gene activation. These discrete state variables depict the extent to which gene expression has deviated from its basal level at any point in time. Transitions between each of the variables’ states are governed by the stochastic processes listed below.

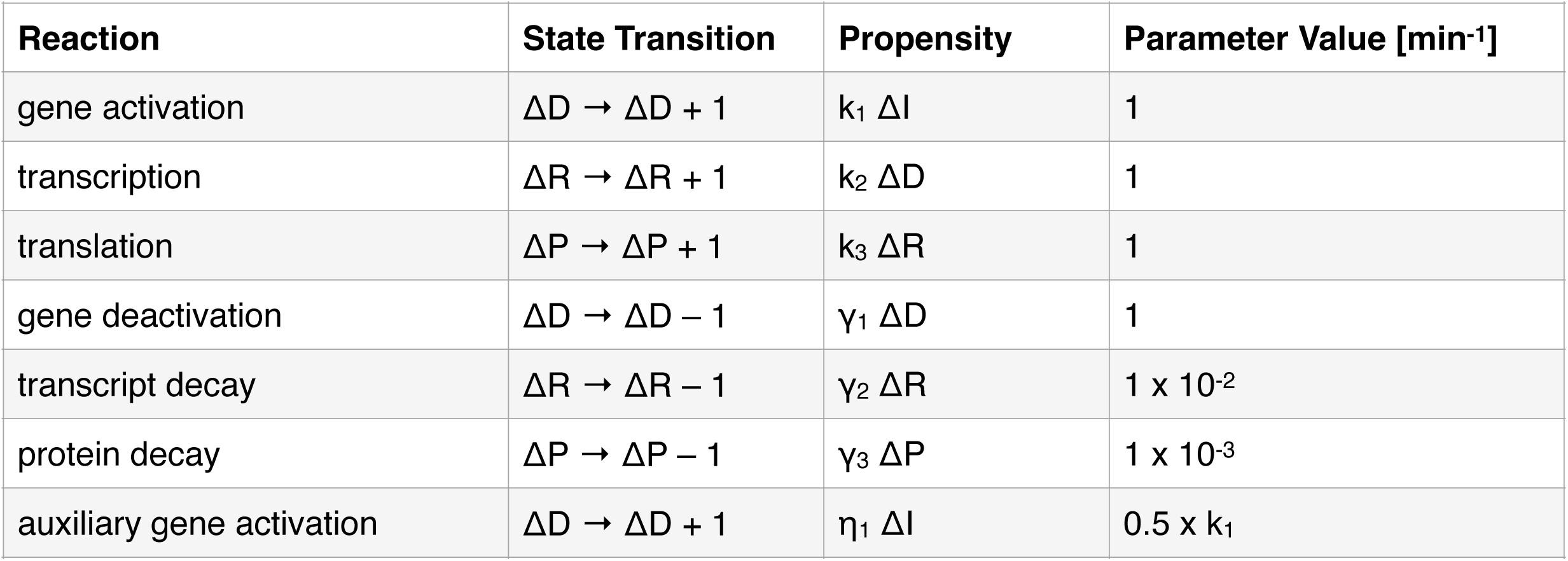
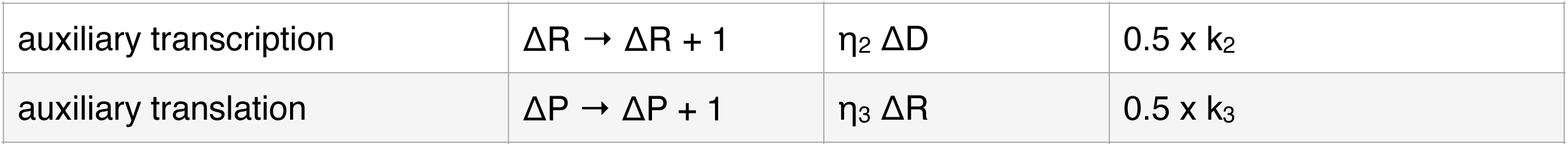

| Reaction | State Transition | Propensity | Parameter Value [$\text{min}^{-1}$] |
| --- | --- | --- | --- |
| gene activation | $\Delta D \rightarrow \Delta D + 1$ | $k_1 \Delta I$ | 1 |
| transcription | $\Delta R \rightarrow \Delta R + 1$ | $k_2 \Delta D$ | 1 |
| translation | $\Delta P \rightarrow \Delta P + 1$ | $k_3 \Delta R$ | 1 |
| gene deactivation | $\Delta D \rightarrow \Delta D - 1$ | $\gamma_1 \Delta D$ | 1 |
| transcript decay | $\Delta R \rightarrow \Delta R - 1$ | $\gamma_2 \Delta R$ | $1 \times 10^{-2}$ |
| protein decay | $\Delta P \rightarrow \Delta P - 1$ | $\gamma_3 \Delta P$ | $1 \times 10^{-3}$ |
| auxiliary gene activation | $\Delta D \rightarrow \Delta D + 1$ | $\eta_1 \Delta I$ | $0.5 \times k_1$ |
| auxiliary transcription | $\Delta R \rightarrow \Delta R + 1$ | $\eta_2 \Delta D$ | $0.5 \times k_2$ |
| auxiliary translation | $\Delta P \rightarrow \Delta P + 1$ | $\eta_3 \Delta R$ | $0.5 \times k_3$ |

In the continuum limit, this model yields a deterministic system of differential equations:

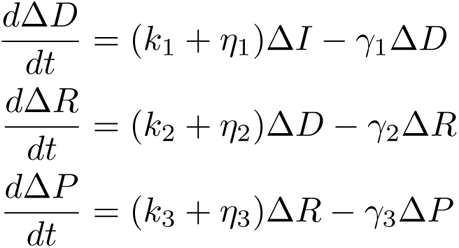

Where *k_i_* are activation, transcription, or translation rate constants, *η_i_* are their auxiliary counterparts, and *γ_i_* are degradation constants. In control parlance, three sequential first-order transfer functions relate input disturbances to deviations in output protein level.

We contend that the Hartman-Grobman theorem holds for the processes under study, but we also recognize that transcriptional and regulatory kinetics are often described using nonlinear kinetics. Therefore, we also considered two nonlinear modeling frameworks, both of which recapitulated the same results.

### Dependence of Model Parameters on Metabolic Conditions

IPC ablation reduces cellular glucose consumption. Presumably this would affect the production and consumption of ATP. Since ATP concentration remains fairly constant when respiration is limited [46], ATP flux (and ATP synthesis) is assumed to decrease. Because transcription, translation, and protein degradation all require ATP turnover, we halved their rate parameters under conditions of reduced glucose consumption. All auxiliary activator strengths were reduced in an equivalent manner to their primary counterparts. That is, rate constants for auxiliary activators driving either transcription or translation were each halved under conditions of reduced glucose consumption. These assumptions are incorporated as changes to the model’s rate parameters as listed below. Conversely, to model the effects of increased ATP consumption we instead increased each of the relevant rate parameters by 50%.

| Parameter | Normal Metabolism | Reduced ATP Consumption | Increased ATP Consumption |
| --- | --- | --- | --- |
| transcription rate constant | $k_2$ | $(0.5)k_2$ | $(1.5)k_2$ |
| translation rate constant | $k_3$ | $(0.5)k_3$ | $(1.5)k_3$ |
| protein decay rate constant | $\gamma_3$ | $(0.5)\gamma_3$ | $(1.5)\gamma_3$ |
| auxiliary transcription activator rate constant | $\eta_2$ | $(0.5)\eta_2$ | $(1.5)\eta_2$ |
| auxiliary translation activator rate constant | $\eta_3$ | $(0.5)\eta_3$ | $(1.5)\eta_3$ |

When modeling the regulation by repressors, we adopted the same assumptions presented in Cassidy et. al. [9]. Specifically, rate constants are assumed to exhibit quadratic dependence on ATP consumption owing to the intermediate transcriptional and translational processes required to express each repressor. This culminates in a 75% reduction in the strength of each repressor under conditions of reduced energy metabolism, and a 225% increase in the strength of each repressor under conditions of elevated energy metabolism.

| Parameter | Normal Metabolism | Reduced ATP Consumption | Increased ATP Consumption |
| --- | --- | --- | --- |
| auxiliary repression of transcription | $\eta_1$ | $(0.25)\eta_1$ | $(2.25)\eta_1$ |
| auxiliary transcript degradation | $\eta_2$ | $(0.25)\eta_2$ | $(2.25)\eta_2$ |
| auxiliary protein degradation | $\eta_3$ | $(0.25)\eta_3$ | $(2.25)\eta_3$ |

### Model Simulations

Default parameter values were based on approximate transcript and protein synthesis and turnover rates for animal cells reported in the literature [47], while gene activation and decay rates were arbitrarily set to a significantly faster timescale. Default strengths for auxiliary activators acting at the gene, transcript, or protein levels were chosen such that ∼60% of simulations failed to reach the threshold protein level under normal conditions when the auxiliary activator was lost. Population-wide expression dynamics were estimated by simulating 5,000 output trajectories in response to a three-hour transient step input to the gene activation rate. Simulations were performed using a custom implementation of the stochastic simulation algorithm (Gillespie, 1977). The algorithm constrains solutions to the set of discrete positive values, consistent with linearization about a basal level of zero gene activity. This simplifying assumption is based on the near-zero basal activities expected in the experimental systems, but is not required to support the conclusions of the model (Figures S2-C and S4-C).

### Evaluation of Error Frequencies and Changes in Expression Dynamics

Gene expression trajectories were simulated both with (full activation) and without (partial activation) auxiliary activators. Protein expression dynamics were compared by evaluating the fraction of partial-activation simulation trajectories that fell below the top 99% of full-activation trajectories at each point in time, *t*. We refer to this under-expression value as *E(t)*. The time point at which the full-activation simulations mean level reached 30% of its maximum value was taken to be the commitment time. Under-expression was averaged across the time course, beginning with the reception of the input and ending at the commitment time, *τ*.

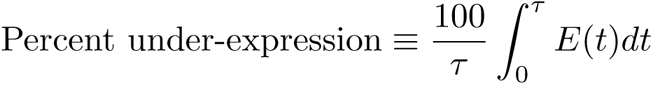

Percent under-expression reflects the net extent to which the expression dynamics differ between the two sets of simulated trajectories.

To estimate the error frequency due to loss of auxiliary activators, the instantaneous error rate was computed by evaluating *E(t)* at the time at which the full-activation simulations mean level reached its maximum amplitude. Because the threshold was set at the bottom 1% of full-activation protein levels, the minimum possible error frequency is one percent. For simplicity we subtracted this percentage point from all reported error frequencies.

### Parameter Variation and Sensitivity to Model Assumptions

We conducted parameter sweeps to confirm the robustness of each computational result. In each sweep, all model parameters were varied across a ten-fold range (± ∼three-fold). We quasi-randomly generated 1,000 such parameter sets, then independently ran four sets of 5,000 simulations for each: 1) full activation with normal metabolism, 2) partial activation with normal metabolism, 3) full activation with reduced metabolism, 4) partial activation with reduced metabolism. Partial-activation systems were assigned a single primary activator for each stage of synthesis. In addition to these primary activators, each full-activation system was assigned an additional set of auxiliary activators whose strengths relative to the primary activators were specified by a free parameter we refer to as *severity*. Error frequencies were evaluated as described above.

Each sweep sampled a seven-dimensional space. Projecting the results of all simulations onto each of the 21 orthogonal 2-D planes revealed that error frequency is greater than 1% for almost all combinations of parameter values (Figure S1A). While it helps illustrate our parameter sweep methodology, the 2-D visualization does not offer sufficient insight into the global trend to justify its complexity. We instead opted for a condensed 1-D projection (Figure S1B), which clearly indicates that loss of auxiliary activators induces an increase in error frequency across a broad parameter range. Auxiliary activator loss also decreases protein levels throughout the time course for the vast majority of parameter sets (Figure S3A).

The difference in error frequency between simulations with normal metabolism and reduced metabolism are shown in Figures S1C, and S2 for all parameter sets. There is a general trend of decreased error frequency with partial activation under reduced energy metabolism conditions. The difference in protein under-expression between simulations with normal versus reduced metabolism are shown for all parameter sets in Figures S3B and S4. Most parameter sets show less under-expression in the absence of auxiliary activators when metabolism is reduced.

### Alternate Models

The number of active sites firing transcription within a cell is limited by gene copy number, but the activated-DNA state in our simple linear model is unbounded. To test whether error frequency suppression persists when an upper bound on gene activity is introduced, we considered a simple two-state transcription model whose deterministic representation is given by:

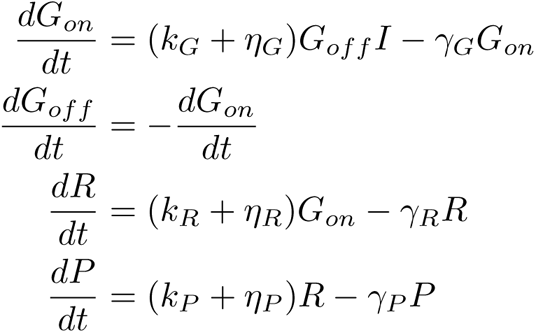

where *G_on_* and *G_off_* are the on- and off-states of a gene; *I*, *R* and *P* are the input, RNA, and protein levels; *k_i_*, γ*_i_*, and η*_i_* are the primary synthesis, decay, and auxiliary synthesis rate constants for species *i*, respectively. Rate parameter dependencies upon metabolic and protein synthesis conditions were analogous to those used in the linear model, and are tabulated below.

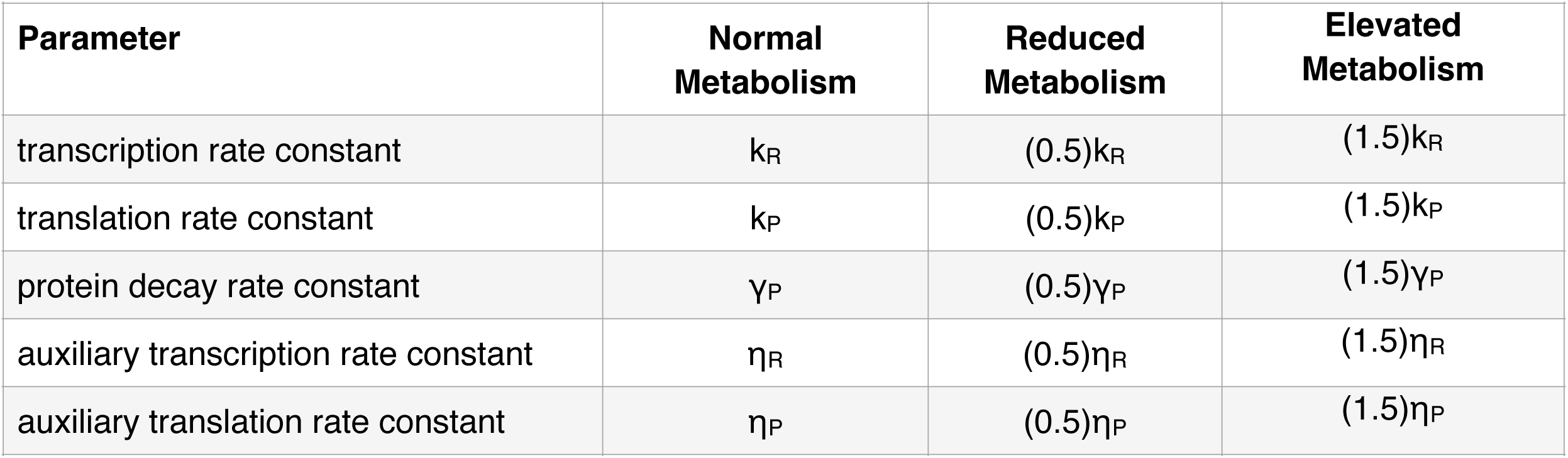

We performed a parameter sweep of this model in which all simulations were initialized as diploid (*G_off_* = 2). Despite the limitation placed on gene activity, error frequency remains elevated under normal growth conditions and partially suppressed when metabolism is reduced (Figure S2A).

Gene expression models also frequently utilize cooperative kinetics in order to capture the nonlinearities and thresholds encountered in transcriptional regulation. We reformulated our gene expression model in terms of Hill kinetics:

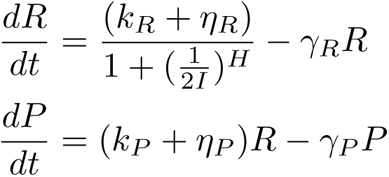

where *I*, *R*, and *P* are the input, RNA, and protein levels; *k_i_*, γ*_i_*, and η*_i_* are the primary synthesis, decay, and auxiliary synthesis rate constants for species *i*; and *H* is a transcriptional Hill coefficient. The stimulus level corresponding to half-maximal transcription rate was fixed at 0.5 because we only consider a binary input signal. Rate parameters were again scaled with metabolic conditions in a manner analogous to the linear model.

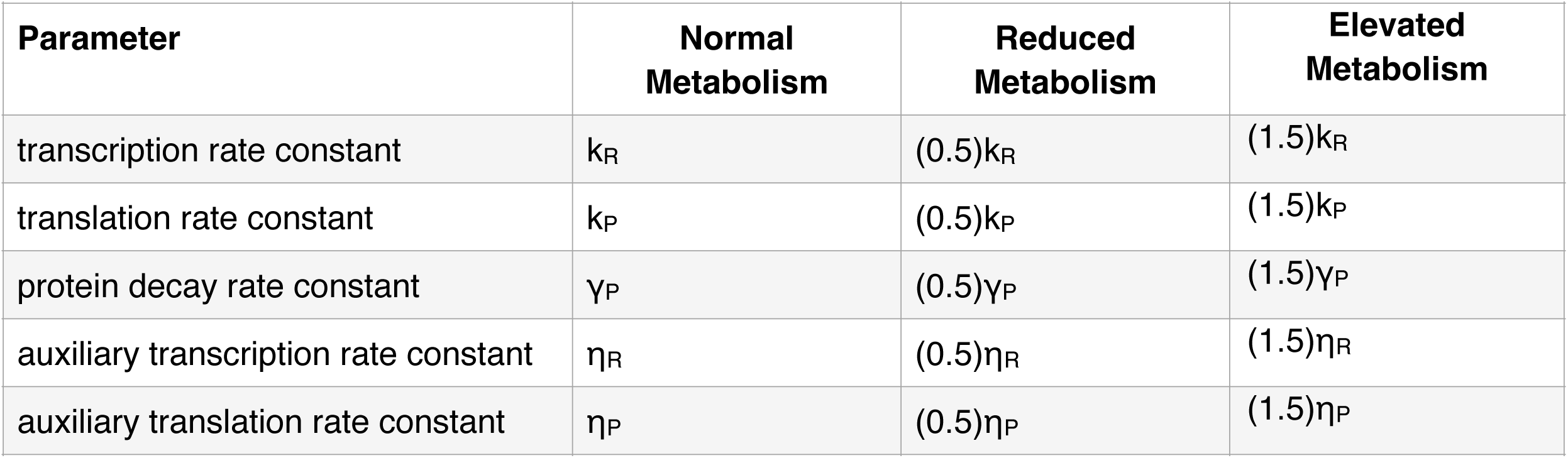

Another parameter sweep revealed that despite the incorporation of cooperative binding kinetics, error frequency remains elevated under normal metabolic conditions and is broadly suppressed when metabolism is reduced (Figure S2B).

### Quantification and Statistics

Confidence intervals for the moving average of Yan-YFP expression were inferred from the 2.5th and 97.5th percentile of 1,000 point estimates of the mean within each moving-average window. Point estimates were generated by bootstrap resampling with replacement of the expression levels within each window. Histogram distributions for the mean value of the *D* metric were calculated from 10,000 point estimates of the mean as generated by bootstrap resampling with replacement of *D* metric measures for each condition. Best-fit Gaussian distributions were fit onto each histogram. There was no exclusion of any data or subjects.

## Data Availability

The experimental datasets generated and analyzed during the current study are available at https://doi.org/10.21985/n2-5bf9-5e21.

Data of all model simulations and figures presented in this manuscript is available at https://doi.org/10.21985/n2-j361-8e86

## Code Availability

All code is publicly available for free.

Modeling and analysis: https://github.com/sbernasek/promoters

The *Silhouette* app used to segment nuclei in eye discs and annotate nuclei in all eye discs is freely available via the macOS App Store.

The *FlyEye* python package used to generate, align, and visualize expression dynamics has been published on GitHub (https://doi.org/10.5281/zenodo.5520468).

The *roughEye* MATLAB package used to segment, correct and analyze lattice disorder in adult compound eyes has been published on GitHub (https://github.com/K-D-Gallagher/roughEye).

## SUPPLEMENTARY FIGURE LEGENDS

**Figure S1. Model expression dynamics are dependent on gene activators - related to Figure 1.** Each of the seven model parameters was varied by one order of magnitude centered around the default value as defined in the STAR Methods. 1,000 such variable parameter sets were generated. Simulations with full and partial activation were performed for each parameter set. **(A)** Protein output was compared between full and partial activation over the entire time course of gene expression. Shown is the frequency in which output reduction occurs with partial activation when compared to full activation for all parameter sets. **(B)** Error frequencies with partial activation were calculated using a threshold set at the lowest 1% of peak protein output from full activation. Shown is a grid of all 21 pairwise combinations of parameter variations. Error frequencies are projected as color heat maps on the 21 squares. Error frequencies are high (light brown) for many combinations of parameter values. **(C)** Distribution of error frequency for all parameter sets under conditions of normal energy metabolism.

**Figure S2. Transient loss of EGFR activity causes lower Yan output - related to Figure 3. (A)** Yan expression dynamics in *Egfr^f24^*/+ heterozygous eyes after incubation in vivo at either 18°C or 26.5°C for 18 hours before fixation and analysis. N refers to number of single cells in which Yan measurements were made. **(B)** Yan expression dynamics in *Egfr^tsla^*/+ heterozygous eyes after incubation in vivo at either 18°C or 26.5°C for 18 hours before fixation and analysis. N refers to number of single cells in which Yan measurements were made. **(C)** Yan expression dynamics in *Egfr^tsla^*/*Egfr^f24^* trans-heterozygous eyes after incubation in vivo at either 18°C or 26.5°C for 18 hours before fixation and analysis. N refers to number of single cells in which Yan measurements were made.

**Figure S3. Representative images of Yan expression in eye discs - related to Figure 3.** All panels show a single optical slice through cell nuclei in the field of view taken of eye discs from animals that were incubated at 26.5°C for 18 hours. Left panels, DAPI channel; middle panels, Yan-YFP channel; right panels, merged channels. **(A)** An *EGFR* wildtype animal with normal unperturbed metabolism. **(B)** An *EGFR* mutant animal with normal unperturbed metabolism. **(C)** An *EGFR* wildtype animal that is also *Dilp2>Rpr*. **(D)** An *EGFR* mutant animal that is also *Dilp2>Rpr*. **(E)** An *EGFR* wildtype animal that is also *GMR>Myc*. **(F)** An *EGFR* mutant animal that is also *GMR>Myc*.

**Figure S4. Workflow for quantitative analysis of disorder in the compound eye - Related to Figure 5. (A)** Sample adults mounted on Blu-Tack. **(B)** Correct configuration of fiber-optic lighting of samples on the Leica microscope stage. **(C)** Raw image of an eye using correct lighting and exposure conditions. **(D)** Example of incorrect configuration of lighting in which extraneous objectives distort the lighting. **(E)** Resulting images with incorrect lighting and exposure are shown. **(F,G)** An image of an eye after segmentation of the ommatidia **(F)** and subsequent erosion of the segmented region of interest **(G)** to minimize distortion due to eye curvature. **(H)** Centroids within segmented ommatidia are interconnected by a Delaunay triangulation. The nearest neighbors of each centroid are identified. **(I)** Distances from each centroid to its nearest neighbors are calculated.

**Figure S5. Brightfield images of adult compound eyes - related to Figure 5.** Shown are representative raw images of eyes for disorder quantitation. Lighting and exposure are adjusted to generate pointillistic reflection of light from ommatidial lenses. **(A)** *Egfr^f24^ / +* wildtype. **(B)** *Egfr^tsla^ / Egfr^f24^* mutant. **(C)** *mir-7^Δ1^ / +* wildtype. **(D)** *mir-7^Δ1^* mutant.

**Figure S6. Density distributions of the mean *D* for EGFR mutants with normal or reduced metabolism - related to Figure 6.** Density distributions of the mean *D* were estimated for ommatidia from wildtype (purple) and *EGFR ts* mutant (green) eyes. All animals were raised at 18°C except for an 18-hour interval as late L3 larvae when they were incubated at 26.5°C. **(A,B)** Eye disorder in animals with normal metabolism. **(C)** Eye disorder in animals with reduced metabolism due to IPC ablation.

