## Supplementary figures and images for "Energy metabolism modulates the regulatory impact of activators on gene expression"

### Supplemental Figures

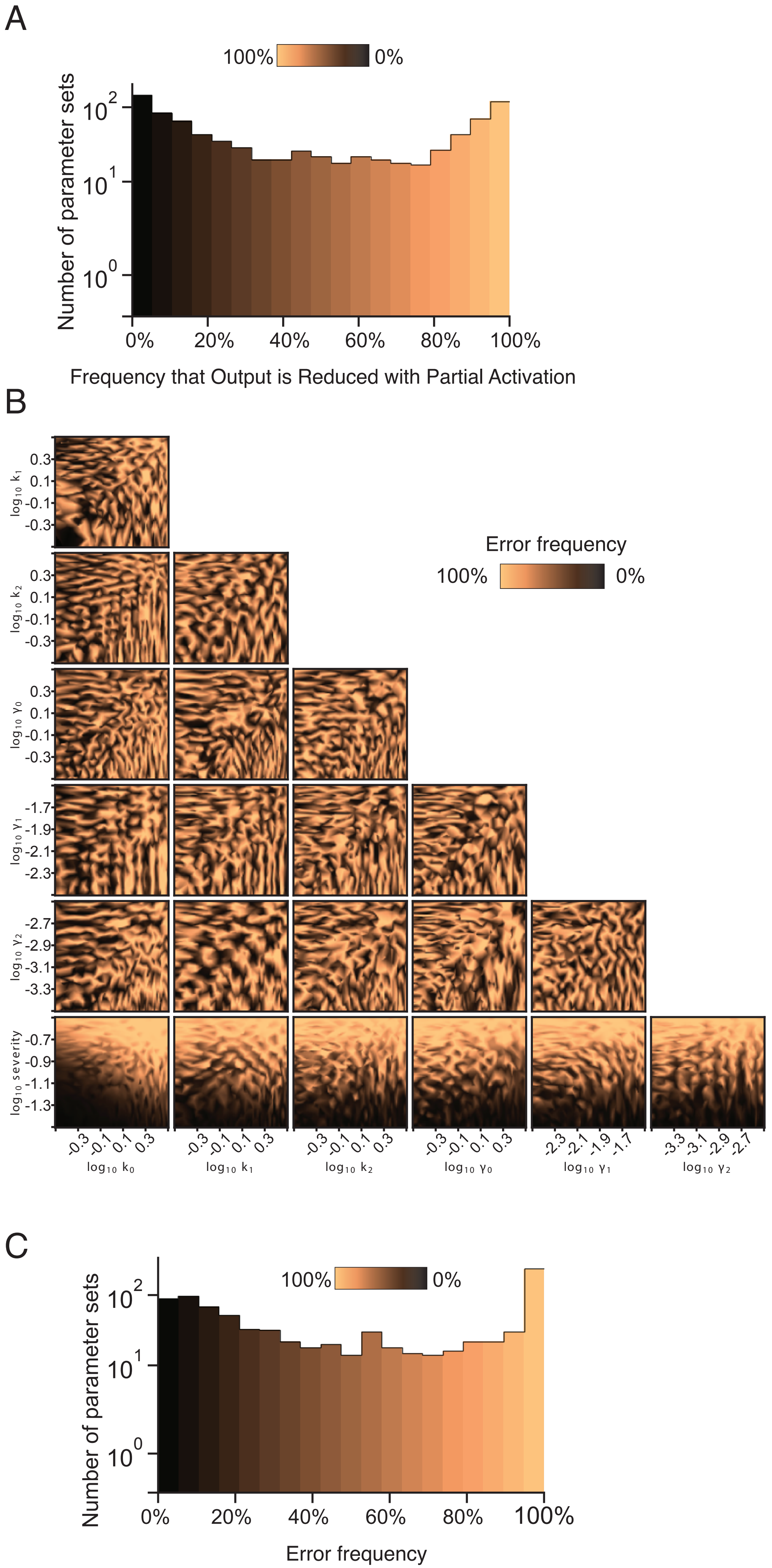

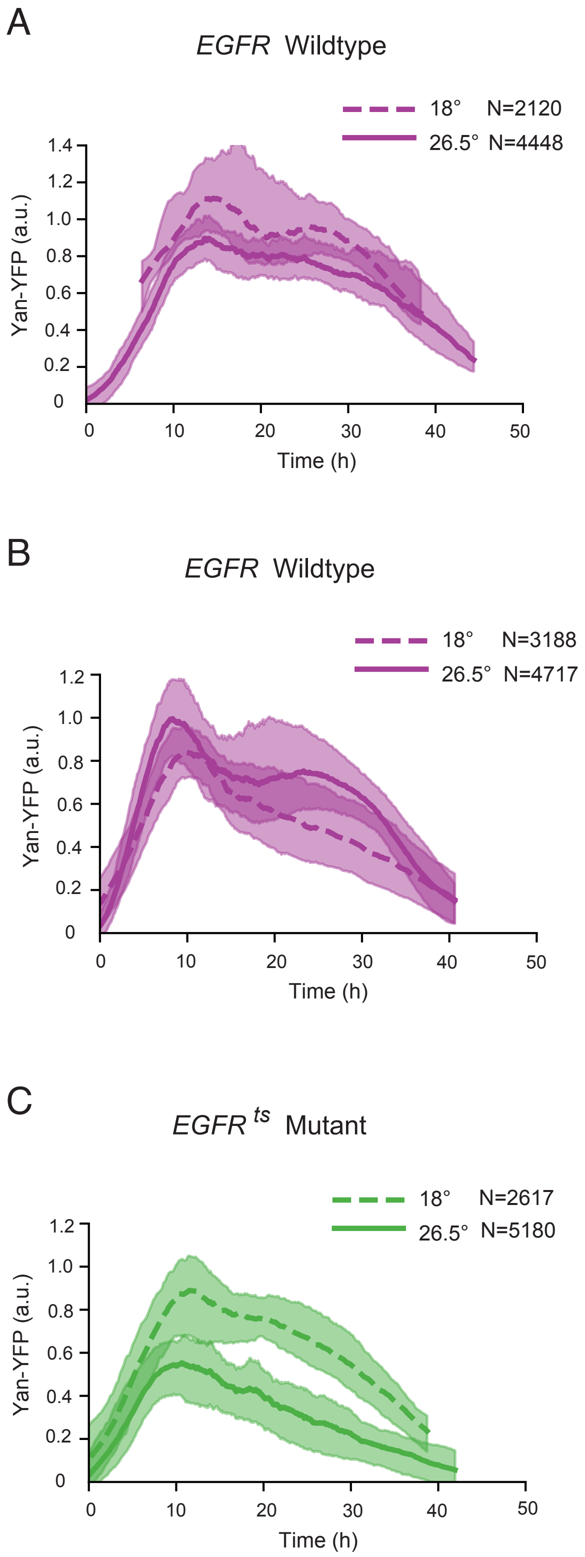

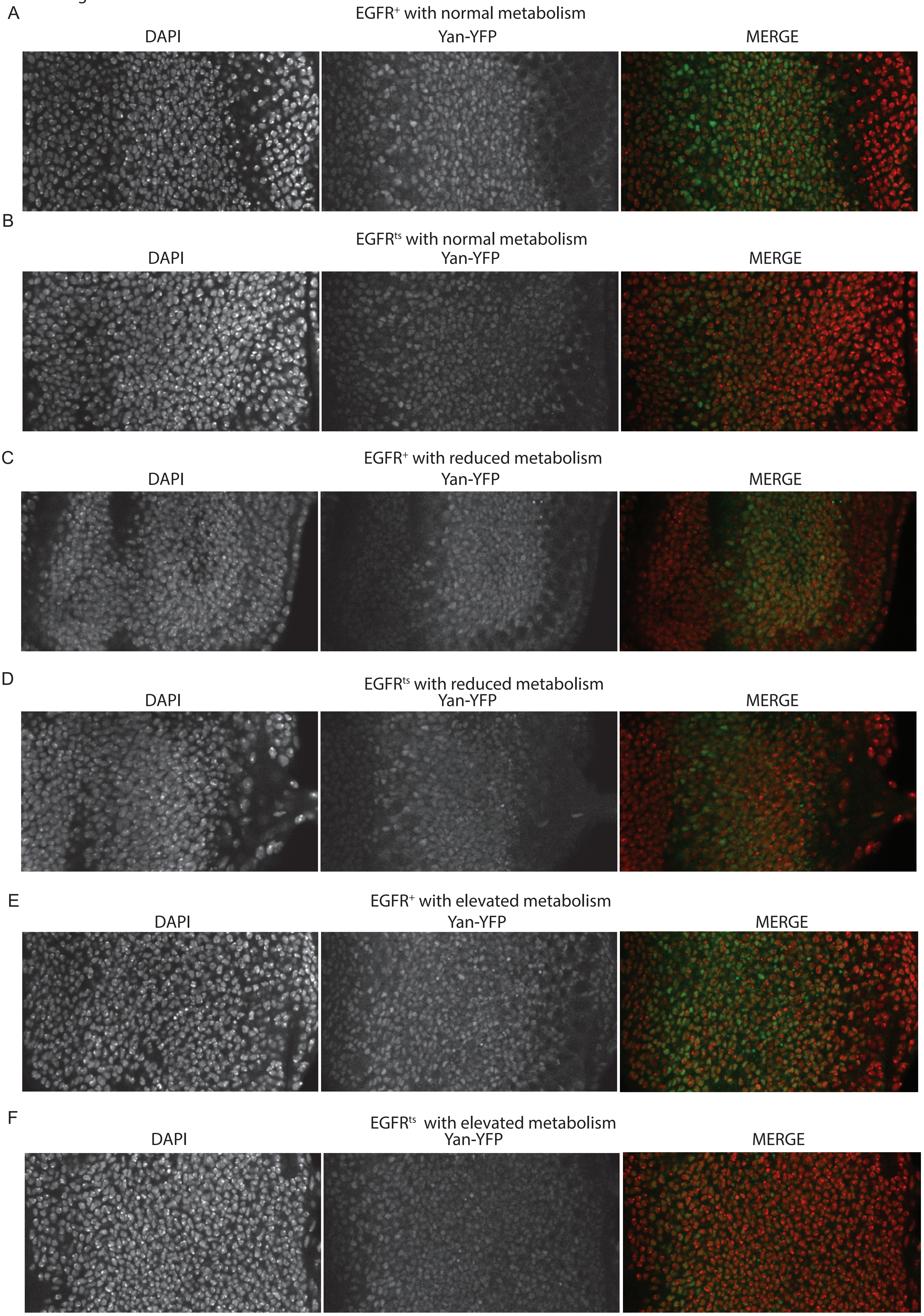

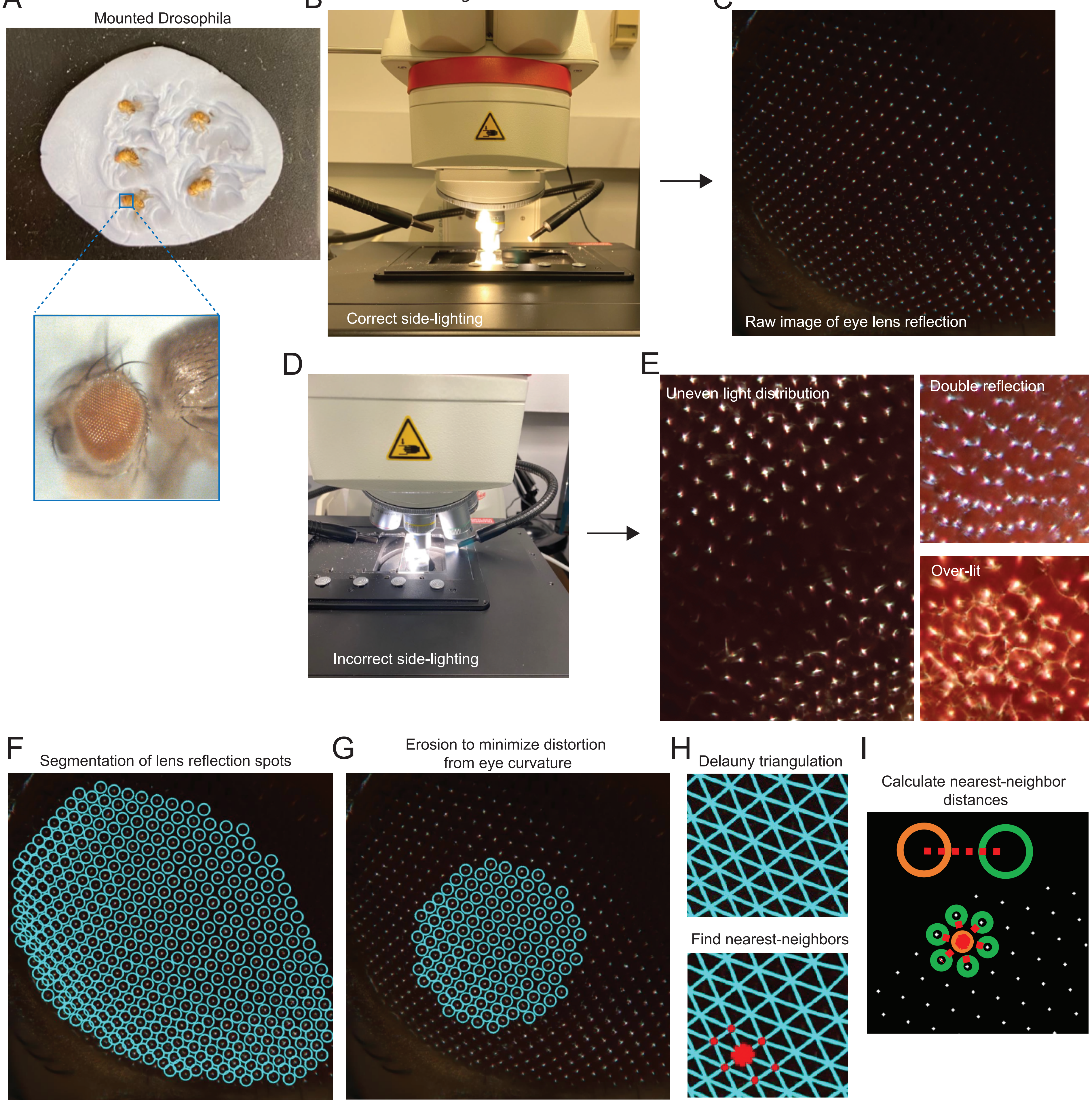

Figure S5

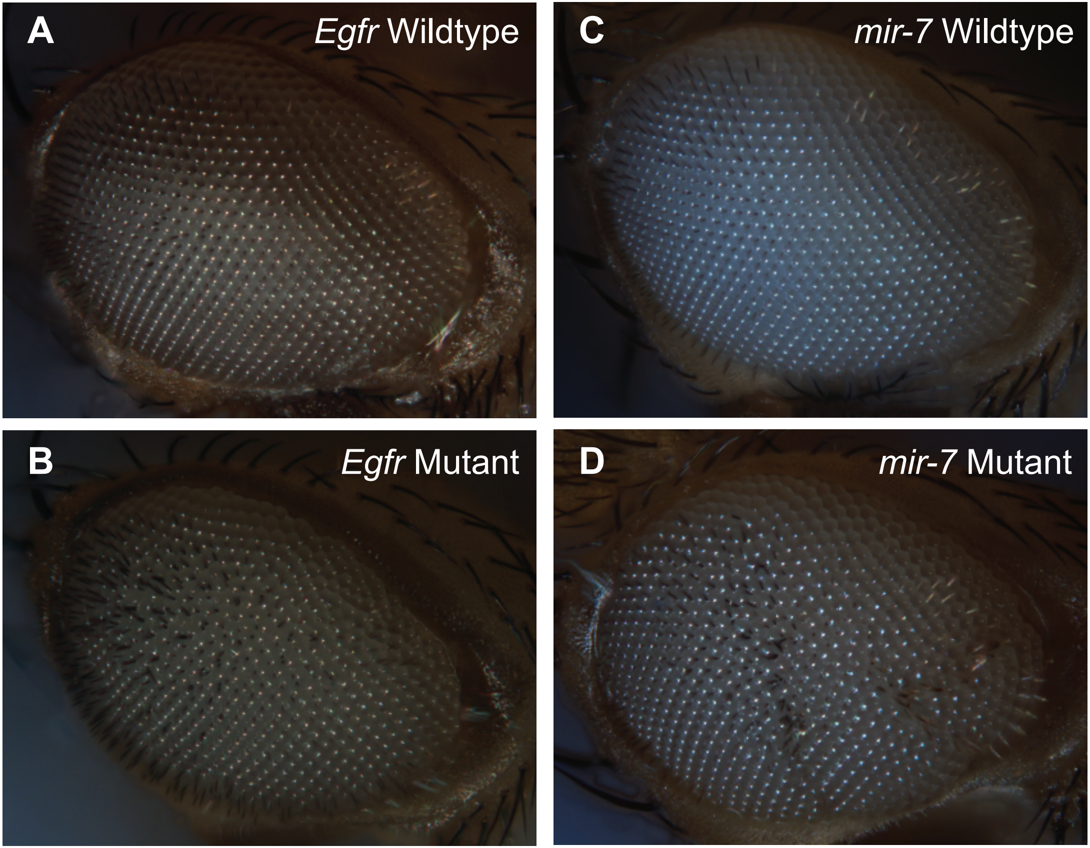

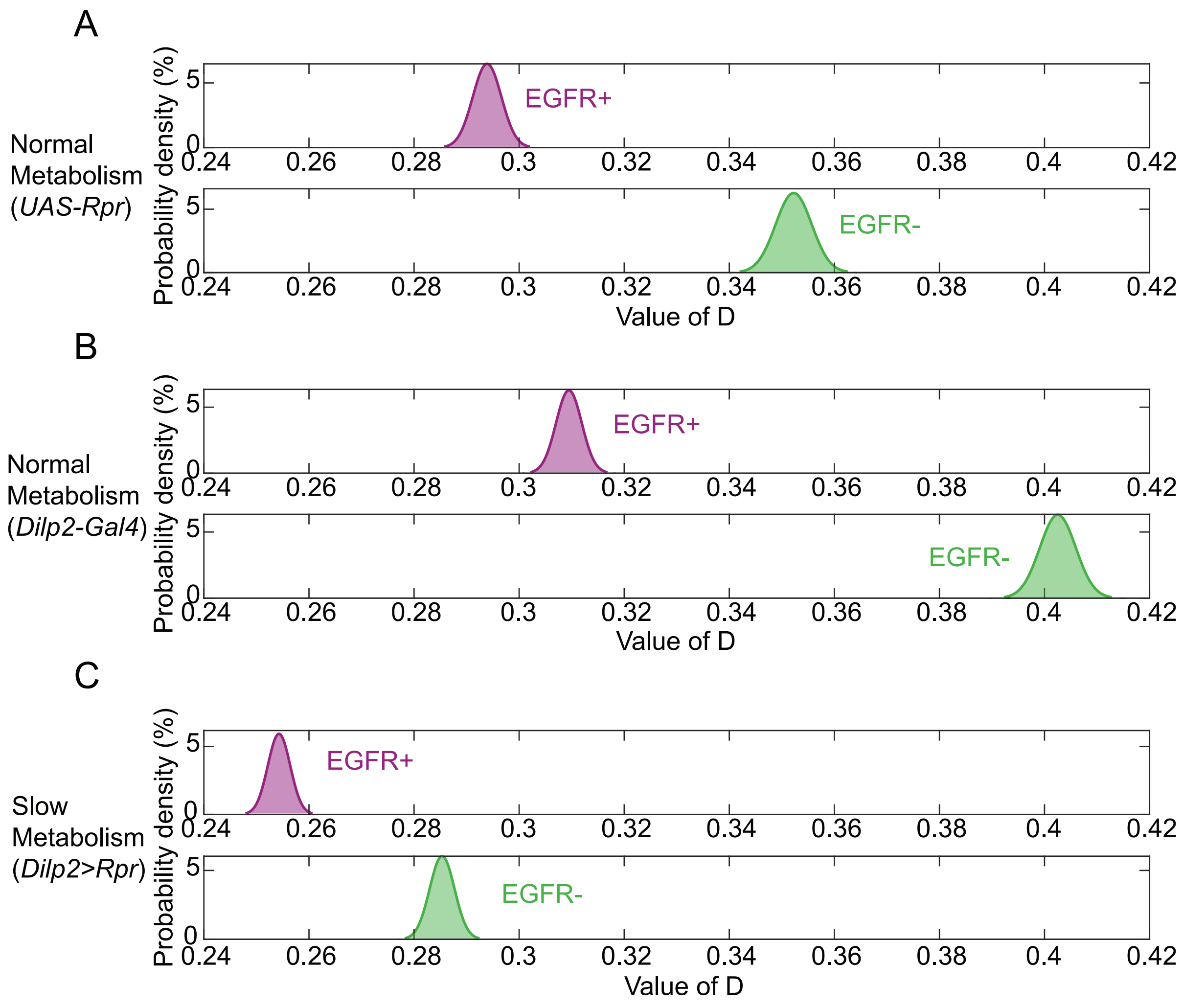
